# Hypoxia-induced SETX links replication stress with the unfolded protein response

**DOI:** 10.1101/2020.05.24.113548

**Authors:** Shaliny Ramachandran, Tiffany Ma, Natalie Ng, Iosifina P. Foskolou, Ming-Shih Hwang, Pedro Victori, Wei-Chen Cheng, Francesca M. Buffa, Katarzyna B. Leszczynska, Natalia Gromak, Ester M. Hammond

## Abstract

The levels of hypoxia associated with resistance to radiotherapy significantly impact cancer patient prognosis. These levels of hypoxia initiate a unique transcriptional response with the rapid activation of numerous transcription factors in a background of global repression of transcription. Here, we show that the biological response to radiobiological hypoxia includes the induction of the DNA/RNA helicase SETX. In the absence of hypoxia-induced SETX, R-loop levels increase, DNA damage accumulates, and DNA replication rates decrease. SETX plays a key role in protecting cells from DNA damage induced during transcription in hypoxia. Importantly, we show that the mechanism of SETX induction is reliant on the PERK/ATF4 arm of the unfolded protein response. These data not only highlight the unique cellular response to radiobiological hypoxia, which includes both a replication stress dependent DNA damage response and an unfolded protein response but uncover a novel link between these two distinct pathways.

## INTRODUCTION

The microenvironment of the majority of solid tumours is characterised by regions of low oxygen (hypoxia), which result from abnormal vasculature and increased oxygen demand. Tumour hypoxia plays a pivotal role in tumourigenesis, and has been associated with increased aggressiveness, metastasis, resistance to radiation therapy, and poor patient prognosis ^1^. The level of hypoxia associated with radiation resistance, radiobiological hypoxia (<0.13% O_2_), is characterised by a rapid induction of replication stress, which is defined as the slowing or stalling of replication forks. We have attributed hypoxia-induced replication stress to decreased nucleotide availability ^2-4^. Previously, we demonstrated that hypoxia-induced replication stress leads to the induction of an ATR- and ATM-dependent DNA damage response (DDR). However, hypoxia-induced replication stress is insufficient to induce ATM-mediated signalling and increased levels of heterochromatic histone marks, most notably H3K9me3 are also required ^5-7^. Importantly, both hypoxia-induced ATR and ATM activity occur in the absence of detectable DNA damage ^8^.

R-loops are 3–stranded nucleic acid structures which are linked to transcription and can contribute to replication stress ^9^. R-loops are comprised of an RNA/DNA hybrid and a displaced single-stranded DNA (ssDNA). R-loops form within the genome at actively transcribed genes preferentially occurring at gene promoters and terminators, but also in response to the loss of RNA biogenesis factors, and at sites of head-on transcription-replication collisions ^10-13^. R-loops regulate gene expression but can also lead to genome instability, which is particularly apparent in cells with deficiencies in R-loop processing and binding factors, including for example BRCA1 ^14-18^. Many different factors including RNase H enzymes, RNA/DNA helicases, topoisomerases, and mRNA biogenesis/processing factors are involved in the processing and resolution of R-loops ^19,20^. The hypoxic environment is known to significantly change the transcriptional network of the cell, suggesting that there may also be an impact on R-loop formation and resolution ^21,22^. DNA repair pathways, including critical components such as BRCA1 and Rad51, have been shown to be repressed through a variety of mechanisms in response to hypoxia ^23-25^. In contrast, a number of transcriptional pathways are induced in radiobiological hypoxia including the hypoxia-inducible factors (HIF1-3), p53 and those initiated by the unfolded protein response (UPR) ^26-28^. The UPR is an ER stress response pathway that leads to global translation inhibition while upregulating selective genes including those involved in protein folding, redox homeostasis and protein degradation ^29^. In contrast to the DDR, which is activated by ssDNA in hypoxia, the UPR is activated by misfolded proteins.

Senataxin (SETX) is a putative RNA/DNA helicase that can facilitate transcription termination and gene transcription of a subset of genes ^12,30-33^. The yeast homolog of SETX, Sen1, was shown to be involved in transcription-coupled repair and protect cells from transcription-associated recombination ^34-36^. In addition, Sen1 has been shown to act at the replication fork and facilitate replication at highly transcribed genes ^37^. SETX has also been linked to R-loops at transcription-replication collisions and to participate in the antiviral response ^36,38^. Mutations in SETX have been associated with two neurodegenerative diseases, ataxia with oculomotor apraxia type 2 (AOA2) and amyotrophic lateral sclerosis type 4 (ALS4) ^39,40^.

The role of R-loops and factors involved in R-loop processing including RNA/DNA helicases have not been investigated in response to hypoxia. We show that in radiobiological hypoxia, in the context of transcriptional and replication stress, R-loops increase. The majority of factors associated with R-loops were repressed with the notable exception of SETX, which was specifically induced in radiobiological hypoxia, prompting us to focus on the role of SETX in hypoxia. We found that the loss of SETX lead to an accumulation of R-loops, and that hypoxia-induced SETX protected cells from replication stress, transcription dependent DNA damage and apoptosis. Most importantly, hypoxia-induced SETX was dependent on the PERK branch of the UPR, providing the first evidence of a link between the UPR and replication stress in hypoxia.

## RESULTS

### SETX is induced in an oxygen-dependent manner

Exposure to hypoxia (<0.1% O_2_) led to the rapid accumulation of RPA foci and pan-nuclear γH2AX in the absence of 53BP1 foci. Importantly, the cells that had pan-nuclear γH2AX also had RPA foci which indicates a replication stress-dependent DDR despite the absence of detectable DNA damage (Figure 1A, B, S1A). The cellular response to radiobiological hypoxia included the rapid repression of global transcription rates as shown by quantification of 5-ethynyluridine (5’EU) incorporation (Figure 1C, D, S1B). In these same conditions (<0.1% O_2_), the transcript levels of a number of R-loop processing factors including RNase H1, RNase H2B, PIF1, DHX9, RTEL1 and AQR, were significantly repressed. In contrast, SETX mRNA was upregulated in hypoxia (<0.1% O_2_) (Figure 1E) and this was evident in a number of cell lines (Figure S1C-E). SETX has not previously been shown to be stress-responsive at the transcriptional level. SETX protein levels were also upregulated in hypoxia in various cell lines including RKO, A549 and HCT116 (Figure 1F-H). To determine oxygen dependency, we compared expression levels in <0.1% O_2_ to 2% O_2_ and found that SETX was specifically induced at <0.1% O_2_ (Figure 1I). To investigate whether this oxygen dependent expression of SETX was evident in cancer patients, we used The Cancer Genome Atlas (TCGA) datasets for colorectal and lung cancers. Here, we compared SETX expression with both, a validated hypoxia signature and with a group of genes that we have previously shown are induced at <0.1% O_2_ but not 2% O_2_ ^26,41^. As we saw no indication of SETX induction in 2% O_2_, we were not surprised to find that SETX expression did not correlate with a hypoxia signature (Figure S1F). However, in agreement with the SETX induction at <0.1% O_2_, SETX expression showed a positive correlation in the three datasets used with the group of p53-target genes specifically induced at <0.1% O_2_ (Figure 1J). Therefore, it seems likely that SETX expression is increased in patient samples from tumours with radiobiological levels of hypoxia, lending further support to our novel finding that SETX is induced in an oxygen-dependent manner.

**Figure 1.**
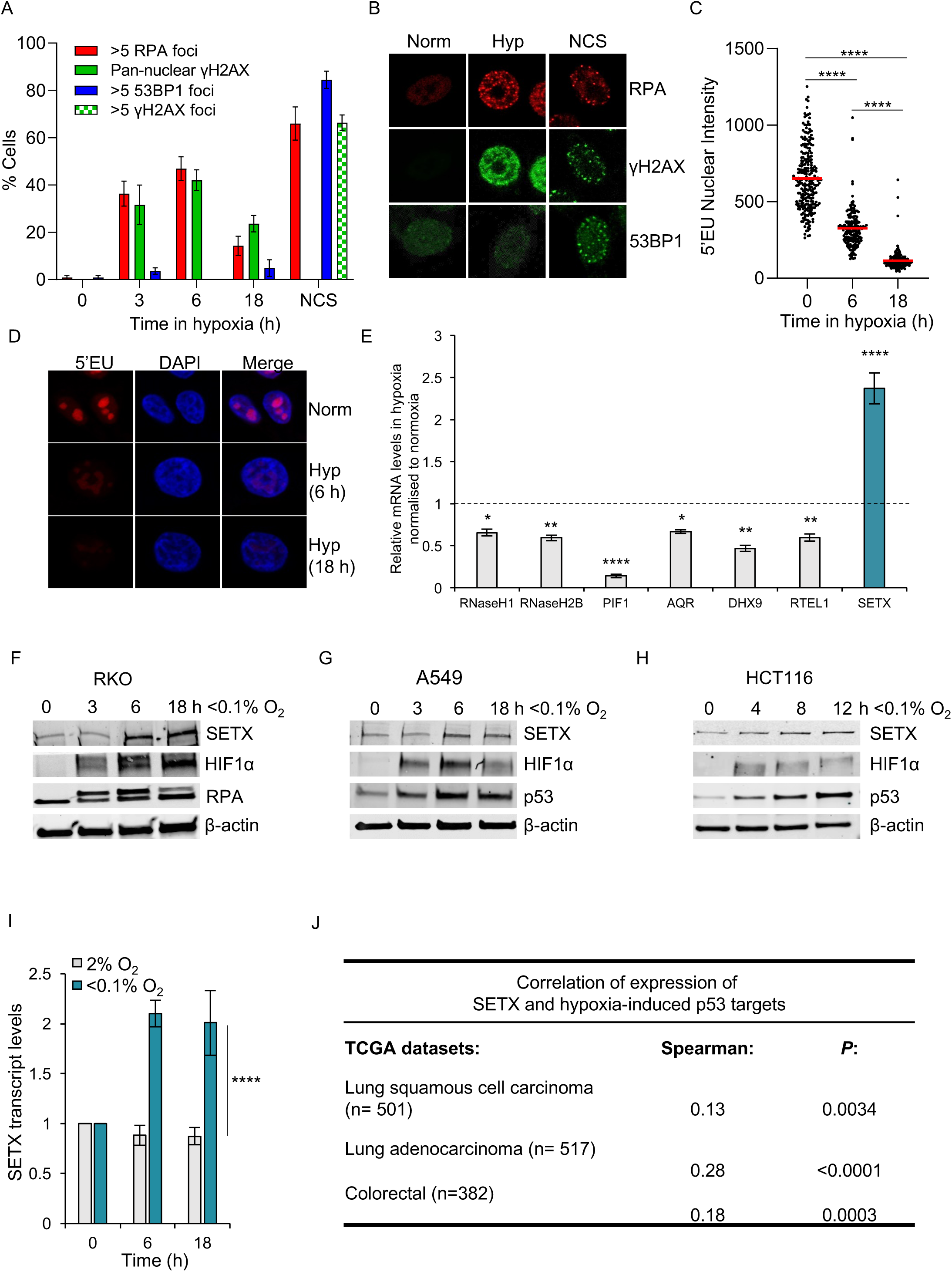
SETX is induced in an oxygen dependent manner. A. RKO cells were exposed to <0.1% O_2_ for 0, 3, 6 and 18 hours or to NCS (200 ng/µL) for 6 hours. Cells were fixed and stained for γH2AX, RPA and 53BP1. A representative plot (of 2 independent) experiments is shown. B. Representative images from A. C. RKO cells were exposed to <0.1% O_2_ for 0, 6 and 18 hours with 5’EU (0.5 mM) added for the final 6 hours. Representative of 3 independent experiments. D. Representative images from C. E. RKO cells were exposed to <0.1% O_2_ for 6 hours followed by RT-qPCR for the mRNAs indicated relative to the normoxic control. The dashed line indicates normoxic expression levels. F. RKO cells were treated for 0, 3, 6 and 18 hours of <0.1% O_2_. The levels of SETX, HIF1α and RPA are shown. β-actin is the loading control. G. A549 cells were treated for 0, 3, 6 and 18 hours of <0.1% O_2_. The levels of SETX, HIF1α and p53 are shown. β-actin is the loading control. H. HCT116 cells at 0, 4, 8 and 12 hours of treatment with <0.1% O_2_. The levels of SETX, HIF1α, p53 and the loading control β-actin, are shown. I. A549 cells were exposed to hypoxia, 2% O_2_ or <0.1% O_2_ for the times indicated and the relative mRNA levels of SETX determined. J. To examine *SETX* expression against tumour-associated hypoxia-induced p53-activity (referred to as hypoxia-induced p53 targets), raw data for each sequenced gene were rescaled to set the median equal to 1, and hypoxic p53 targets were determined by quantifying the median expression of 6 p53 target genes associated with hypoxia-induced p53 activity (encoding *BTG2, CYFIP2, INPP5D, KANK3, PHLDA3* and *SULF2*) ^26^. Correlations and statistical significance were determined by calculating Spearman’s rho rank correlation coefficient (r) and two-tailed *P* value using Hmisc package in RStudio. (A-I) Data from three independent experiments (n=3), mean ± standard error of the mean (SEM) are displayed unless otherwise indicated.

### SETX alleviates R-loops and replication stress in hypoxia

To determine the impact of loss of SETX in hypoxia on R-loops, we used the S9.6 antibody, which specifically recognises RNA/DNA hybrids. R-loop levels increased in hypoxia in A549 and RKO cells and this increase was ablated when the cells were treated with RNase H, an endonuclease that degrades the RNA component of R-loops (Figure 2A, B and S2A). SETX depletion led to an increase in the levels of R-loops in both normoxia and hypoxia, and again, this increase was ablated with RNase H treatment (Figure 2C and S2B-F). R-loops can contribute to replication stress, so we asked if SETX had a role to play in replication in hypoxia. Depletion of SETX led to a significant increase in stalled replication forks in normoxic conditions (Figure 2D). However, no changes were detected in hypoxic conditions, likely because hypoxia alone induced high levels of stalled forks (Figure 2E). SETX depletion decreased replication rates in both normoxia and hypoxia, however, this was more significant in the hypoxic conditions (Figure 2F and S2G, H).

**Figure 2.**
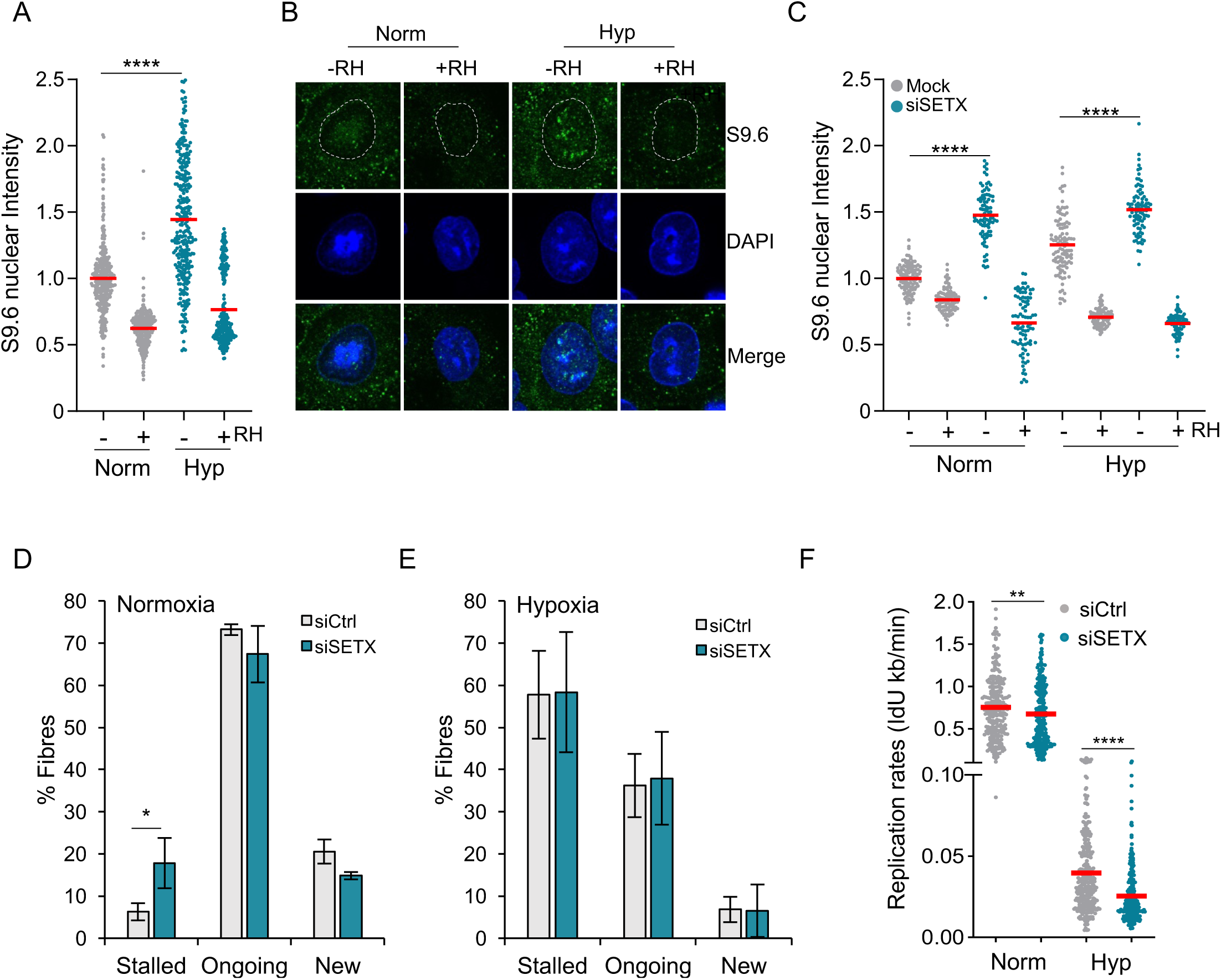
SETX alleviates R-loops levels and replication stress in hypoxia. A. A549 cells were exposed to 21% O_2_ or <0.1% O_2_ for 12 hours, fixed and stained with the S9.6 antibody and DAPI. Where indicated, coverslips were treated with RNase H prior to staining. B. Representative images from A. Dashed white line indicates nuclear outline based on DAPI stain that was used to measure the nuclear S9.6 intensity. C. A549 cells that were either mock transfected (grey) or transfected with SETX siRNA (green), were exposed to 21% O_2_ or <0.1% O_2_ for 12 hours. Cells were fixed and stained with the S9.6 antibody and DAPI. Where indicated, coverslips were treated with RNase H prior to staining. n=1 D. RKO cells transfected with control siRNA (grey) or SETX siRNA (green), were subjected to DNA fibre assays at 21% O_2_ (Normoxia) and the different types of replication structures were quantified. E. RKO cells transfected with control siRNA or SETX siRNA, were subjected to DNA fibre assays at <0.1% O_2_ (Hypoxia) and the different types of replication structures were quantified. F. Replication rates, as determined by IdU incorporation rates, of RKO cells transfected with control siRNA versus SETX siRNA exposed to 21% O_2_ (Norm) or <0.1% O_2_ (Hyp) for 6 hours. (A-F) Data from three independent experiments (n=3), mean ± SEM are displayed unless otherwise indicated.

### SETX protects hypoxic cells from transcription associated DNA damage and apoptosis

SETX has been suggested to affect RNA polymerase II (Pol II) binding at certain genes and to facilitate transcriptional termination of a number of genes ^12,32,42^. Under normoxic conditions SETX-knock-down lead to a significant decrease in 5’EU incorporation suggesting that SETX plays a role in global transcription. In hypoxia where transcription is repressed, there was a modest, further decrease in 5’EU incorporation upon SETX depletion (Figure 3A and S3A). A previous study identified changes in the expression of 36 genes in response to SETX depletion, and 60 genes changing in a SETX-dependent manner in response to viral infection ^38^. We carried out RNA-sequencing (RNA-seq) and found that SETX depletion only affected a small subset of genes (17 increasing and 48 decreasing) in normoxia, and of these genes, 5 were previously identified as regulated in a SETX-dependent manner. In contrast, SETX depletion had a much higher impact in hypoxia causing 341 genes to increase and 256 genes to decrease in expression (Figure 3B). These results were validated by RT-qPCR (Figure S3B). We investigated whether the differentially expressed genes shared any common features by gene ontology analysis and found that approximately 20% of genes upregulated in SETX depleted hypoxic cells were involved in rRNA processing, nucleolus and ribosome biogenesis (Figure S3C). We also investigated whether the differentially expressed genes shared any other common features. Hypoxia-induced genes that had increased expression in SETX depleted cells had a slightly shorter gene length compared to the hypoxia-induced genes that were unaffected or had decreased expression upon SETX depletion (Figure S4A). There was no evidence for the genes differentially expressed upon SETX depletion that are also hypoxia inducible or repressive, having similar number of exons, number of G4 quadruple structures or GC percent (Figure S4B-D).

**Figure 3.**
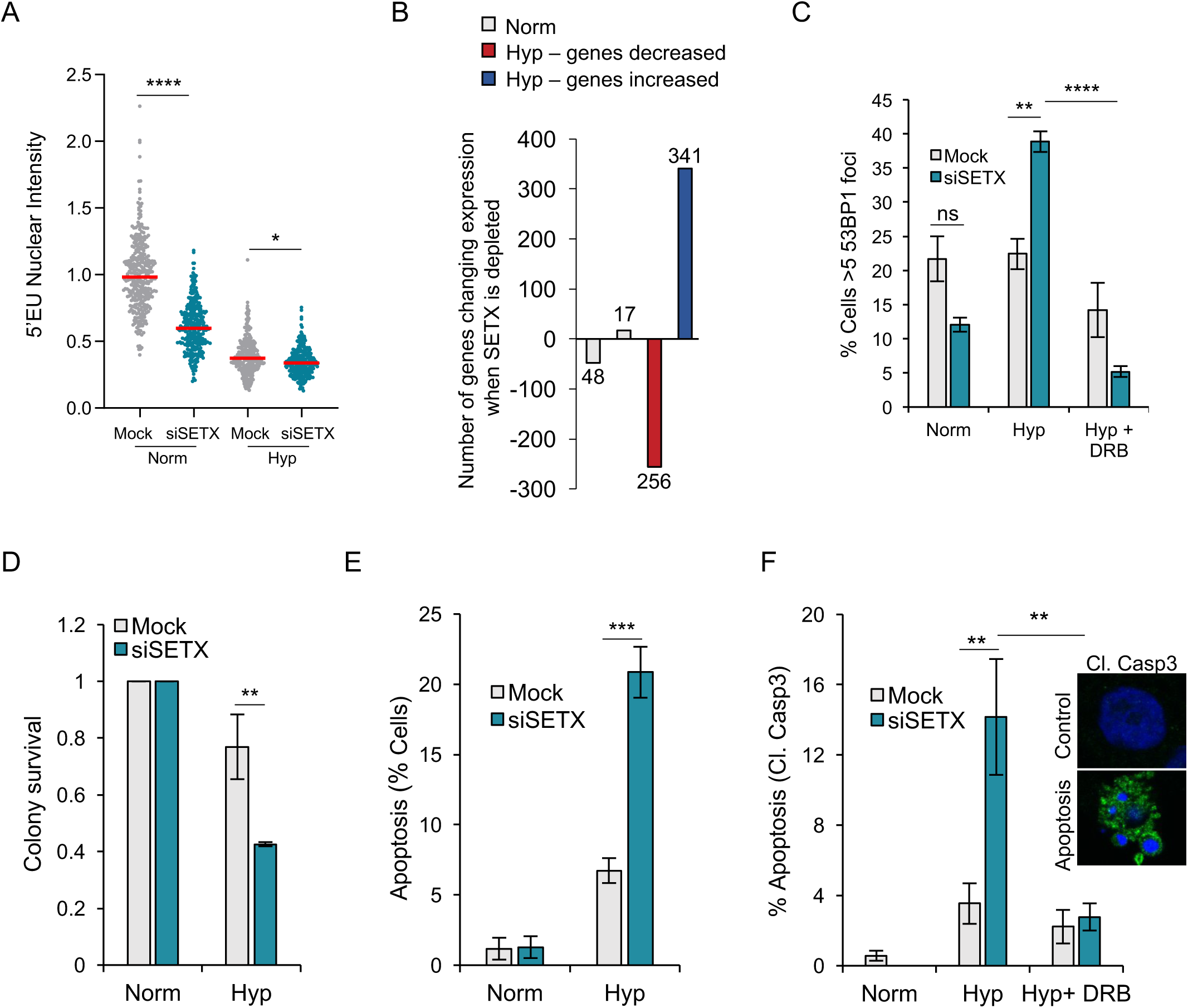
SETX protects hypoxic cells from transcription associated DNA damage and apoptosis. A. RKO control cells or SETX depleted cells were exposed to 21% and <0.1% O_2_ for 6 hours in the presence of 5’EU. Cells were fixed, stained and quantified for 5’EU (0.5 mM) incorporation. B. RNA-seq results comparing RKO cells treated with control siRNA or SETX-siRNA in 21% O_2_ (Norm, grey) and <0.1% O_2_ (Hyp). Genes with decreased expression upon SETX depletion in hypoxia shown in red, and genes with increased expression upon SETX depletion in hypoxia shown in blue. RNA-seq was carried out in 3 biological replicates (n=3). C. Percentage of cells displaying 53BP1 foci (>5 per nucleus) in RKO cells with either mock treatment or SETX siRNA. Cells were exposed to 21% O_2_ (Norm), 6 hours of <0.1% O_2_ (Hyp) or pre-treated with DRB (100 µM) followed by 6 hours of <0.1% O_2_ (Hyp + DRB). D. Colony survival of A549 cells with either mock treatment or SETX siRNA after 0 and 18 hours of hypoxia (Hyp) (<0.1% O_2_) treatment. E. A549 cells were treated with SETX siRNA or mock and exposed to 21% O_2_ (Norm) or <0.1% O_2_ (Hyp) for 24 hours. The percentage of cells undergoing apoptosis as determined by DAPI staining is shown. F. A549 cells were mock treated or treated with SETX siRNA and exposed to 21% O_2_ (Norm), <0.1% O_2_ (Hyp) for 24 hours or pre-treated with DRB (100 µM) followed by 24 hours of <0.1% O_2_ (Hyp + DRB). The percentage of cells positive for cleaved caspase 3 is shown. Representative images of a normal and apoptotic cell are included. (A-F) Data from three independent experiments (n=3), mean ± SEM are displayed unless otherwise indicated.

Given that unresolved R-loops have been associated with DNA damage, we asked whether depleting SETX would lead to DNA damage in hypoxia. As expected, there was no increase in DNA damage in hypoxia but upon SETX depletion there was a significant increase in DNA damage as determined by the increase in cells with 53BP1 foci (Figure 3C and S5A). To test whether the DNA damage observed in the SETX depleted hypoxic cells was linked to transcription we treated cells with dichloro-beta-D-ribofuranosylbenzimidazole (DRB), which inhibits Pol II-mediated transcription. DRB rescued the DNA damage observed in the SETX depleted hypoxic cells suggesting that the damage observed in SETX depleted hypoxic cells was transcription dependent (Figure 3C). We confirmed that DRB did not rescue DNA damage non-specifically by demonstrating that DRB did not rescue Adriamycin-induced DNA damage (Figure S5B). It should be noted that by inhibiting transcription, DRB led to a decrease in R-loop levels, suggesting that SETX may be protecting the hypoxic cells from co-transcriptional R-loop associated DNA damage (Figure S5C-E). The damage observed in SETX-depleted hypoxic cells is independent of cell cycle occurring in both S-phase and non-S-phase cells, supporting the hypothesis that the damage could be the result of transcription dependent R-loops (Figure S5F, G). DNA damage, if unrepaired, can lead to increased apoptosis and reduced survival. In hypoxia, SETX depletion led to decreased colony survival when compared to control cells, suggesting that SETX function contributes to hypoxic cell viability (Figure 3D). SETX depletion also led to a significant increase in apoptosis compared to the control cells specifically in hypoxia and again, this was found to be transcription dependent as the addition of DRB rescued the apoptosis induced by SETX loss (Figure 3E, F). Taken together, SETX protects hypoxic cells from transcription linked DNA damage and apoptosis.

### SETX upregulation is dependent on the UPR

To investigate the mechanism of SETX induction in hypoxia, A549 cells were used as these showed the most robust transcriptional induction of SETX in hypoxia. SETX mRNA induction occurred at <0.1% O_2_, and multiple candidate transcription factors are active at this oxygen level, including HIF, p53 and those involved in the UPR (ATF3, ATF4, ATF5 and CHOP) (Figure S6A, B). Given that SETX was not induced at 2% O_2_ where HIF is active, and SETX expression did not correlate with a hypoxia signature, it seemed unlikely that SETX induction was dependent on HIF. However, to formally test this, we depleted HIF1α and asked whether this affected SETX induction in hypoxia. As expected SETX induction in hypoxia was not affected by HIF1α siRNA in contrast to the control gene VEGF, confirming that SETX is not induced by HIF1α in hypoxia (Figure S6C, D). Next, we found that SETX induction in hypoxia was not affected by p53 siRNA when compared to control cells, suggesting that p53 signalling does not contribute to SETX induction in hypoxia (Figure 4A, S6E). In support of this conclusion, we and others have found no evidence of a functional p53 response element in the SETX promoter ^43^. Next, we investigated whether SETX induction could be linked to the UPR. Predicted binding sites were identified for the UPR transcription factor ATF4 in the GeneHancer identifier GH09J132349 corresponding to the SETX promoter region. We depleted ATF4 using siRNA and demonstrated that ATF4 is responsible for the induction of SETX in hypoxia (Figure 4B, S7A). The transcript levels of the ATF4 target CHOP were tested in parallel to confirm ATF4 knock-down (Figure S7B). Chromatin immunoprecipitation qPCR was used to confirm that ATF4 is responsible for SETX induction in hypoxia. ATF4 was enriched at the SETX promoter specifically in hypoxia (Figure 4C). ATF4 enrichment at the CHOP gene was used as a positive control (Figure S7C). To further investigate whether SETX is a UPR target, we exposed cells to a range of stresses including tunicamycin and thapsigargin, both of which are known to induce the UPR, and found that both led to a significant increase in SETX mRNA similar to the induction observed in hypoxia (<0.1% O_2_) (Figure 4D, S7D). Consistent with this, SETX was found to be upregulated in a RNA sequencing dataset upon tunicamycin addition although this was not specifically reported ^44^. As expected, we also demonstrated that SETX expression was not induced with hypoxia mimetics desferrioxamine (DFO) and 2% O_2_, which lead to HIF stabilisation (Figure 4D, S7E). Importantly, hydroxyurea did not affect SETX transcript levels demonstrating that hypoxia-mediated induction of SETX is UPR dependent and distinct from replication stress (Figure 4D). Thapsigargin and tunicamycin induced SETX but did not lead to replication stress as observed by an absence of RPA phosphorylation (Figure 4E, S7F).

**Figure 4.**
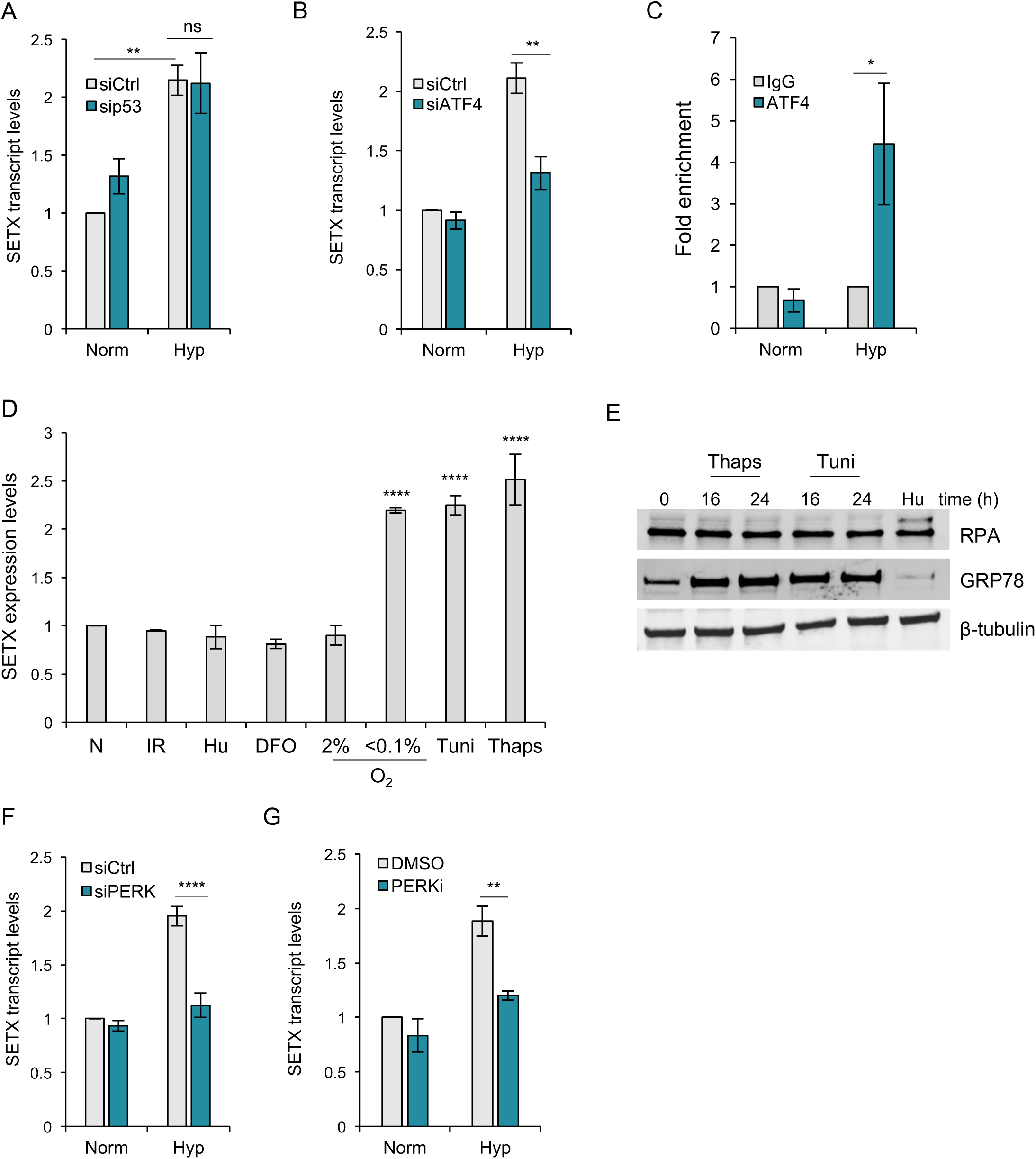
SETX upregulation is dependent on the PERK branch of the UPR. A. A549 cells were treated with either control siRNA or p53 siRNA and exposed to 21% O_2_ (Norm) or <0.1% O_2_ (Hyp) for 6 hours. SETX mRNA levels were determined by RT-qPCR. B. A549 cells were treated with either control siRNA or ATF4 siRNA and exposed to 21% O_2_ (Norm) or <0.1% O_2_ (Hyp) for 16 hours. SETX mRNA levels were determined by RT-qPCR. C. A549 cells were exposed to 21% O_2_ (Norm) or <0.1% O_2_ (Hyp) for 4 hours, and ChIP-qPCR experiments were performed to determine fold enrichment of ATF4 at the SETX promoter relative to IgG control. D. SETX mRNA levels in non-treated A549 cells (N), cells exposed to irradiation (5 Gy) followed by a 1 hour recovery, Hu (1 mM), DFO (100 µM), hypoxia (2% O_2_ or < 0.1% O_2_), tunicamycin (Tuni, 5 µg/mL) or thapsigargin (Thaps, 2 µM) for 16 hours. E. A549 cells were exposed to Thaps (2 µM) or Tuni (5 µg/mL) for the times indicated, or to Hu (2 mM, 8 h) followed by western blotting. F. A549 cells were treated with either control siRNA or PERK siRNA and exposed to 21% O_2_ (Norm) or <0.1% O_2_ (Hyp) for 16 hours. SETX mRNA levels were determined by RT-qPCR. n=4 G. A549 cells were treated with either DMSO or PERKi (10 μM) and exposed to 21% O_2_ (Norm) or <0.1% O_2_ (Hyp) for 6 hours. SETX mRNA levels were determined by RT-qPCR. n=4 (A-G) Data from three independent experiments (n=3), mean ± SEM are displayed unless otherwise indicated.

As signalling to ATF4 in hypoxia is primarily mediated by the protein kinase R (PKR)-like endoplasmic reticulum kinase (PERK), we hypothesised that SETX induction in hypoxia would also be PERK dependent ^45,46^. Depleting PERK abrogated the hypoxia-mediated induction of SETX (Figure 4F, S7G). To further validate our finding, we used a PERK inhibitor GSK2606414 (PERKi). Pre-treating cells with PERKi led to a significant decrease in SETX mRNA levels in hypoxia as compared to the control cells (Figure 4G, S7H). We also verified that pre-treating cells with PERKi abrogated SETX induction observed upon thapsigargin treatment, again demonstrating that SETX induction during the UPR is downstream of PERK (Figure S7I, J).

### The UPR is linked with replication stress in hypoxia

Since SETX depletion led to an increase in R-loops, we hypothesised that PERK depletion would have a similar effect. Both, siRNA mediated PERK depletion and pharmacological inhibition of PERK led to an increase in R-loop levels in hypoxia suggesting that, like SETX, PERK signalling negatively regulates R-loops in hypoxia (Figure 5A and S8A, B). PERK depletion or inhibition also partially rescued the global transcriptional repression observed in hypoxia (Figure 5B, S8C). As SETX depletion led to an increase in transcription dependent DNA damage, we asked whether PERK inhibition showed a similar effect. Akin to SETX depletion, PERK inhibition led to an accumulation of DNA damage which was more significant under hypoxic conditions and dependent on transcription (Figure 5C). Additionally, both PERK inhibition and depletion led to increased DDR signalling in hypoxia, specifically increased phosphorylation of p53 and RPA (Figure 5D). PERK inhibition was confirmed by the absence in the electrophoretic mobility shift of PERK on western blot (Figure 5D). Increased RPA phosphorylation with PERK inhibition would indicate a possible increase in replication stress. The percentage of cells experiencing replication stress in hypoxia did not change with PERK depletion, which is expected since hypoxic cells in S phase of cell cycle have replication stress due to the lack of dNTPs (Figure S8D). However, there was an observable increase in the number of RPA foci per cell upon PERK depletion or inhibition in hypoxia suggesting that hypoxic cells with PERK depletion or inhibition may have increased replication stress (Figure 5E, F and S8E). Finally, we tested whether PERK inhibition led to increased replication stress in hypoxia by DNA fibre assay. Inhibition of PERK led to decreased replication rates in both normoxia and hypoxia, suggesting that PERK signalling acts to alleviate replication stress (Figure 5G). Taken together, our data suggests that in radiobiological hypoxia, the PERK branch of the UPR signals through SETX and potentially other unidentified factors to reduce R-loops, replication stress, DNA damage and apoptosis (Figure 5H).

**Figure 5.**
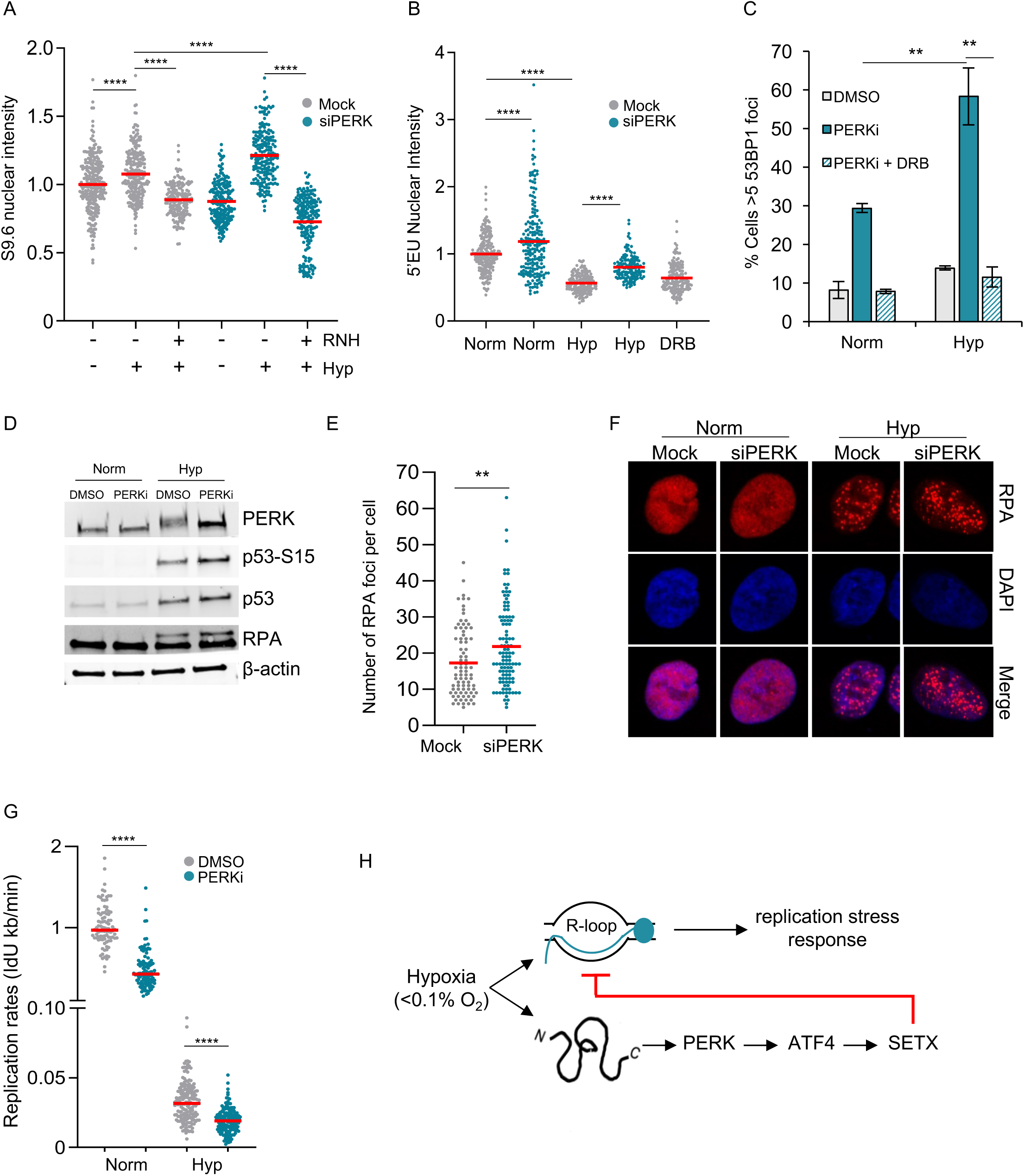
The UPR is linked with replication stress in hypoxia. A. A549 cells that were either mock transfected (grey) or transfected with PERK siRNA (green), were exposed to 21% O_2_ or <0.1% O_2_ for 6 hours. Cells were fixed and stained with the S9.6 antibody and DAPI. Where indicated, coverslips were treated with RNase H (RNH) prior to staining. B. A549 control cells or PERK depleted cells were exposed to 21% and <0.1% O_2_ for 6 hours, 5’EU (0.5 mM) was added for the final hour of treatment. DRB was used as the positive control. Cells were fixed, stained and quantified for 5’EU incorporation. C. The percentage of cells displaying 53BP1 foci (>5 per nucleus) as determined by immunofluorescence, in A549 cells exposed to 21% O_2_ (Norm) or 6 hours of <0.1% O_2_ (Hyp). Cells were pre-treated with DMSO, PERKi (10 µM), or DRB (100 µM) and PERKi (10 µM) (PERKi + DRB). n=3 D. A549 cells were treated with DMSO or PERKi (10 µM) and exposed to normoxia or hypoxia (<0.1% O_2_) for 6 hours followed by western blotting as indicated. E. A549 cells with either mock treatment or PERK siRNA were exposed to hypoxia (<0.1% O_2_) for 6 hours, cells were fixed and stained for RPA by immunofluorescence assays. The number of RPA foci per cell was quantified. F. Representative images from E. G. Replication rates, as determined by IdU incorporation rates, of A549 cells treated with DMSO or PERKi (10 µM) and exposed to 21% O_2_ (Norm) or <0.1% O_2_ (Hyp) for 6 hours. Over 90 fibres were measured for each normoxic sample and over 150 fibres for each hypoxic sample, n=1. H. Hypoxia leads to the activation of the UPR and replication stress. In addition to a shortage of nucleotides, R-loops accumulate in hypoxia and contribute to replication stress. The PERK branch of the UPR signals through ATF4 to increase SETX expression in hypoxia. Hypoxia-induced SETX reduces R-loop accumulation and subsequent replication stress-mediated signalling. (A-G) Data from two independent experiments (n=2), mean ± SEM are displayed unless otherwise indicated.

## DISCUSSION

Our study demonstrates that in radiobiological hypoxia R-loops accumulate and the RNA/DNA helicase SETX is induced. Hypoxia-induced SETX alleviates replication stress and protects hypoxic cells from transcription-associated DNA damage and apoptosis. Importantly, SETX induction was dependent on the PERK branch of the UPR, the first evidence of a link between the UPR and the replication stress dependent DDR in hypoxia. Furthermore, PERK inhibition increased R loops and replication stress in hypoxia, uncovering an entirely novel aspect of PERK function in hypoxia.

Although this is the first link between the UPR and replication stress in hypoxia, a previous report showed that in response to a short exposure (1 hour) of thapsigargin or DTT, PERK inhibits replication fork progression, and origin firing through a Claspin/Chk1 pathway ^47^. In contrast, we found that PERK signalling supports replication fork progression in hypoxia, suggesting that chemically induced UPR signalling may have different outcomes to the hypoxia induced UPR ^48^. Alternatively, PERK signalling in response to short periods of the UPR activation may differ from extended UPR-mediated signalling.

While the DDR and UPR are both active in radiobiological hypoxia, the DDR is restricted to S phase cells ^5^. Here, we show for the first time that SETX is responsive to the UPR. The PERK dependency of SETX induction ensures that cells in all phases of the cell cycle increase SETX levels in radiobiological hypoxia, which may be important for protecting cells from co-transcriptional R-loops which are prone to DNA damage in all phases of the cell cycle.

Previous reports have demonstrated that an accumulation of R-loops can lead to an increase in DNA damage. However, radiobiological hypoxia does not lead to an accumulation of DNA damage even after extended periods of time, suggesting that the hypoxia-induced R-loops are distinct to those induced by other genotoxic stresses such as camptothecin and G-quadruplex ligands ^49,50^. One mechanism by which R-loops lead to DNA damage involves the nucleotide excision repair (NER) DNA repair pathway ^51-53^. DNA repair pathways including NER are repressed in hypoxia, which potentially could protect hypoxic cells from R-loop dependent DNA damage particularly after chronic exposures of hypoxia where repression of repair is more evident ^54,55^. We have shown that depleting SETX in hypoxia leads to an increase in DNA damage. The damage in SETX depleted hypoxic cells is transcription dependent and cell cycle independent, supporting the hypothesis that the role of hypoxia induced SETX is to resolve the co-transcriptional R-loops that are prone to DNA damage and therefore reduce genomic instability.

SETX has been suggested as a potential tumour suppressor from a screen using exome and transcriptome sequencing ^56^. Additionally, akin to characterised tumour suppressors, SETX expression was down-regulated in a number of cancers compared to normal tissue controls ^40^. However, AOA2 patients do not present with increased cancer susceptibility. While SETX loss can lead to genome instability, it is possible redundancy with other R-loop resolution mechanisms protect genome stability. In this study, SETX depletion increased R-loops, replication stress, DNA damage and apoptosis in hypoxia suggesting that SETX may promote hypoxic tumour growth, and hence be a potential therapeutic target. It is possible that in the hypoxic microenvironment in which multiple pathways including DNA repair and potentially R-loop resolution factors are repressed, loss of SETX function would not be readily compensated by other mechanisms.

Given that our study showed that inhibiting PERK led to increased R-loops, DNA damage and replication stress in hypoxia, PERK could therefore be a potential therapeutic target. Targeting PERK in cancer has been controversial as PERK has demonstrated both tumour promoting and suppressive activities depending on the type of cancer and intensity of stress ^57^. However, in hypoxia, PERK supports tumour survival and growth, therefore suggesting PERK as a potential therapeutic target specifically for hypoxic tumours ^58,59^. Since PERK inhibition also increased replication stress in hypoxic conditions, we postulate that using PERK inhibitors in conjunction with replication stress inducers such as ATR inhibitors may be beneficial in solid tumours.

## Supporting information

All SI

## ACKNOWLEDGEMENTS

Thank you to Monica Olcina and Kienan Savage for helpful comments on the manuscript. SR, KBL, PV and MH were supported by a CRUK grant C5255/A23755 (awarded to EMH). NN was supported by an MRC studentship (MC_ST_U16007). IPF was supported by CRUK Oxford Centre Prize DPhil Studentship C38302/A12981. NG was supported by a Royal Society University Research fellowship. W-CC was funded by CRUK grant 23969 (awarded to FMB).

## AUTHOR CONTRIBUTIONS

SR designed, conducted and interpreted the experiments and wrote the paper, TM, NN, IPF and MH conducted experiments, PV, W-CC, and FMB carried out the RNA-seq bioinformatics, KBL performed the TCGA analysis and advised on experiments, NG advised on the project, EMH conceived the project, designed and interpreted experiments and wrote the paper. All authors commented on the manuscript.

## METHODS

### Cell lines and reagents

RKO, HCT116 and A549 (ATCC) were grown in DMEM supplemented with 10% FBS and cultured in humidified incubators at 37 °C and 5% CO_2_. Cells were routinely tested for mycoplasma and found to be negative. For plasmid transfections, JetPrime (Polyplus transfection) was used. siRNA transfections were performed using Dharma-FECT-1 (Thermo Fisher Scientific), sequences are available in the SI. siSETX-A siRNA was used for Figures 2D-F, 3B and Supplementary Figures 2D-G, 3, 4, 5A, G. siSETX-B siRNA was used for Figures 2C, 3A, 3C-F and Supplementary Figures 2B, C and 5F. AllStars negative control siRNA (Qiagen) and ON-TARGETplus non-targeting pool (Dharmacon) were used as control siRNA. Drugs used were neocarzinostatin (NCS), DRB, Hydroxyurea, DFO, Adriamycin (Sigma), tunicamycin and thapsigargin (MP Biomedicals), and PERK inhibitor GSK2606414 (Calbiochem). Irradiations were performed using a Gamma Service GSR D1 Cs137 irradiator.

### Hypoxia Treatment

Bactron II and BactronEZ anaerobic chambers (Shel Lab) were used for <0.1% O_2_. An M35 hypoxia work station (Don Whitley Ltd) was used for 2% O_2_. All experiments at <0.1% O_2_ were plated in glass dishes, except for colony survival assays, which were plated in 6 well plates. All hypoxic treatments were harvested within the chamber using equilibrated buffers. Oxygen tensions were confirmed with the Oxylite probe (Oxford Optronix).

### Immunoblotting

Cells were harvested in UTB lysis buffer (9 M Urea, 75 mM Tris-HCl pH 7.5, 0.15 M β-mercaptoethanol). Blots were visualised using the Odyssey Infrared imaging system. Antibodies used were SETX, KAP1 (Bethyl); GRP78, HIF1α, (BD Biosciences); ATM-S1981, ATM (Abcam); RPA, KAP1-S824, p53-S15 (Cell Signaling); p53, β-actin, RNase H1 (Santa Cruz); and γH2AX, H2AX (Millipore).

### RT-qPCR

RNA was extracted using TRI reagent (Sigma), treated with DNase I (NEB) and cDNA prepared using the Verso cDNA synthesis kit (Thermo Scientific). SYBR Green PCR master mix (Applied Biosystems) and 7500 FAST Real-Time PCR machine (Applied Biosystems) were used. The ΔΔCt method was used to determine relative mRNA fold change with 18S as the reference gene. Primer sequences can be found in Table S1.

### Chromatin immunoprecipitation (ChIP) - qPCR

Treated cells were fixed with 1% formaldehyde and quenched in 125 mM glycine. Cells were lysed with SDS lysis buffer (0.5% SDS, 10 mM EDTA, 50 mM tris pH 8.1). After sonication with Diagenode Bioruptor, samples were precleared with protein A agarose beads and subsequently incubated with ATF4 antibody (Cell Signaling) overnight. Antibody-antigen complexes were then pulled down with agarose beads and washed with the following buffers – low salt wash buffer (0.1% SDS, 1% Triton X-100, 2 mM EDTA, 20 mM Tris pH 8.1, 150 mM NaCl), high salt wash (0.1% SDS, 1% Triton X-100, 2 mM EDTA, 20 mM Tris pH 8.1, 500 mM NaCl), LiCl wash buffer (1% Igepal, 10 mM Tris pH 8.1, 250 mM LiCl, 1 mM EDTA, 1% sodium deoxycholate), TE wash buffer (10 mM Tris pH 8.0, 1 mM EDTA). DNA was eluted with 0.1M NaHCO3 and 1% SDS, cross-linking reversed and treated with proteinase K. DNA was purified with Qiagen PCR purification kits and qPCR was carried out using the following primers: SETX For: CTCAGGTGTCTCAGCGGATG, SETX Rev: CGCATTGTTCGCAAGACCTA, CHOP For: AAGAGGCTCACGACCGACTA and CHOP Rev: ATGATGCAATGTTTGGCAAC. The amount of immunoprecipitated material was calculated as the fold enrichment over IgG control.

### Immunofluorescence and confocal microscopy

Cells were fixed in 4% (w/v) paraformaldehyde in PBS, permeabilised in 1% Triton-X-100 in PBS and blocked with 2% (w/v) BSA in 0.1% Tween-20 in PBS. Primary antibodies 53BP1 (Novus Biologicals), RPA (Cell Signaling), cleaved caspase 3 (Cell Signaling) and BrdU (BD Biosciences) were used. For BrdU staining, cells were exposed to BrdU (20 µM) for the final 1 hour in normxoia and 6 hours in hypoxia, after permeabilization cells were denatured in 2 N HCl for 30 min at 37°C, and blocked with 4% (w/v) BSA in 0.05% Tween-20 in PBS before incubation with the antibody. For 5’EU staining, cells were incubated with 5’ethynyluridine (EU) (0.5 mM) (Jena Bioscience) and Click-iT Alexa Fluor 647 labelling kit (Thermofisher) was used to measure nascent transcription. For EdU staining, cells were incubated with EdU (10 µM) and Click-iT Alexa Fluor 647 labelling kit (Thermofisher) was used to measure replication. S9.6 staining protocol was as previously described ^10^. Coverslips were treated with RNase A (Fermentas) for 1 hour at 37°C and where indicated with RNase H (NEB) for 24 hours at 37°C prior to blocking. Coverslips were then incubated with 1:100 diluted S9.6 antibody (Kerafast) for 36 hours at 4°C. Cells were mounted and the entire nucleus imaged using LSM780 or LSM710 confocal microscope (Carl Zeiss Microscopy Ltd). The 5’EU and S9.6 mean nuclear intensity signal was determined using ImageJ plugin/algorithm kindly provided by Dr Kienan Savage, Queen’s University Belfast ^60^. A minimum of 100 cells were included per treatment.

### RNA-seq

RNA was extracted using TRI reagent (Sigma) and treated with DNase I (NEB). The samples were purified using the RNeasy MinElute Clean-up kit (Qiagen) and purity confirmed using a Bioanalyzer. The mRNA fraction was selected for reverse transcription to cDNA. The cDNA generated was end-repaired, A-tailed and adapter-ligated. cDNA libraries were size selected, multiplexed, tested for quality and subjected to paired end sequencing in one lane of a flow cell.

### RNA-seq differential expression analysis

The analyses were run through an established RNA-seq pipeline by the Buffa lab. The technical adapters in the pair-end raw reads were trimmed out by trimmomatic-0.32 and the clear reads were aligned to human genome version GRCh38 release 82 using tophat v2.0.13. The mapped reads from each gene were counted and normalised to fragments per kilobase of transcript per million (FPKM) using Cuffdiff-2.2.1. Fragments from PCR duplication were removed by Picard 1.124. The significance of fold changes of genes among samples were estimated by nonparametric method Rank Product 3.3, a package in R ^61-66^.

### Correlation of SETX expression with TCGA RNA-seq data

RNA-seq data (RNA Seq V2 RSEM) for 382 colorectal adenocarcinoma tumours, 501 lung squamous cell carcinomas and 517 lung adenocarcinomas were downloaded from the TCGA project (accessed through cBioportal: http://www.cbioportal.org/ on the 12^th^ April 2017). To examine *SETX* expression against hypoxia, a validated hypoxia metagene signature was used ^41^. To examine *SETX* expression against tumour-associated, hypoxia-induced p53-activity (referred to as hypoxic p53 targets in the figure), raw data for each sequenced gene were rescaled to set the median equal to 1, and hypoxic p53 targets were determined by quantifying the median expression of 6 p53 target genes associated with hypoxia-induced p53 activity (encoding *BTG2, CYFIP2, INPP5D, KANK3, PHLDA3* and *SULF2*) ^26^.

### DNA fibre analysis

DNA fibre assays were carried out as described ^67^. For normoxic (21% O_2_) treatments, cells were pulse-labelled with CldU (25 mM) for 20 min, washed once with fresh media, followed by the second label IdU (250 mM) for 20 min. For hypoxic (<0.1% O_2_) treatments, cells were incubated in hypoxia for 3 hours before being pulse-labelled with CldU (25 mM) for 2 hours, washed once with fresh media, then followed by IdU (250 mM) for 1 hour. Cells were mounted (Invitrogen) and imaged using LSM780 confocal microscope (Carl Zeiss Microscopy Ltd). Replication rates were calculated as V_ldU_ (kb/min) = [(x ^*^ 0.132 µm) ^*^ 2.59 kb / µm] / t (min), where x = length of IdU.

### Statistical analysis

Statistical tests were performed using GraphPad Prism software (GraphPad Software Inc.) and include one-way ANOVA, two-way ANOVA, 2-tailed Student’s *t* test and Mann-Whitney U tests. ns = non-significant, * = p ≤ 0.05, ** = p ≤ 0.01, *** = p ≤ 0.001, **** = p ≤ 0.0001.

