## Supplementary material for "Hypoxia-induced SETX links replication stress with the unfolded protein response": All SI

Contents

| | | |
|---|---|---|
| SI Tables | Page | 2 |
| SI Methods | Page | 3 |
| SI Figure legends | Page | 4 |
| SI References | Page | 10 |
| SI Figures | Page | 11 |

Table S1 – Primer sequences used

| GENE | FORWARD | REVERSE |
|---|---|---|
| 18S | GCCCGAAGCGTTTACTTTGA | TCCATTATTCCTAGCTGCGGTATC |
| SETX | CTT CAT CCT CGG ACA TTT GAG | TTA ATA ATG GCA CCA CGC TTC |
| RNase H1 | CCTGTACTTACTGGTGTGGA | CCGTGTGAAAGACGCATCTG |
| RNase H2B | GCATCTTGTTGCTGAAACTTC | TCCTTGTCAGTGGAAGCTTG |
| PIF1 | CGAGCCTAGCACAGAAGCC | CCCAGGATTCGCTTTAGCAG |
| AQR | GGCGGCATTAGCTGAAACTG | CAGCTGTAAGCCCATCCACA |
| DHX9 | GCCAATTTCTGGCCAAAGCA | CGAGGCTCAATGGGGAGTTT |
| RTEL1 | GGACTTGGTCAAGAGCGGAA | AGTCAGGTCAAAGGATGCCG |
| TCF19 | GGGCGGTGATCTCTACACCT | ACCTCTTGGGAGTCGGACAT |
| GHITM | CGTCCTTTCGATGTTGCGTC | TTCTTCACAACAGGGGAGGC |
| FAM222A | CCAAGGAGGGCTTCGTCAG | CAGCTCCAGGCTCTTGCTC |
| ZNF367 | TTTCCACGGGAGAAATCGCT | GGTGTGAAGACGCTGATGTG |
| MMP1 | CATGCTTTTCAACCAGGCCC | GGGTACATCAAAGCCCCGAT |
| JUN | GTGCCGAAAAAGGAAGCTGG | CTGCGTTAGCATGAGTTGGC |
| TM4SF | TACACCTTTGCCAGCACTGA | TTGCTGTTGGTGAGAGCAGC |
| GDF15 | GTTGCACTCCGAAGACTCCA | GAGAGATACGCAGGTGCAGG |
| CHOP | GGAGCATCAGTCCCCCACTT | TGTGGGATTGAGGGTCACA |
| VEGFA | CTACCTCCACCATGCCAAGT | CTCGATTGGATGGCAGTAGC |
| ATF4 | TGACCTGGAAACCATGCCAG | AATGATCTGGAGTGGAGGAC |
| PHLDA3 | GCGCCACATCTACTTCACG | CACAAGCCAGAGGGAACAAC |

Table S2 - siRNA sequences used

| siRNA | Sequence |
|---|---|
| SETX-A | GCACGUCAGUCAUGCGUAA,<br>GCAAUAAGCUCAUCCUAGU,<br>GCUCAACUCUCCAAAUAGA,<br>UAGCACAGGUUGUUAAUCA |
| SETX-B | CCAAUUGCUCCUUUCAGGUGUUUGA |
| HIF1α | CUGAUGACCAGCAACUUGAdTdT |
| p53 | GUAAUCUACUGGGACGGAA |
| PERK | CCAAUGGGAUAGUGACGAA,<br>GGUAGGAUCUGAUGAAUUU,<br>GCAAUUAGCCUUAAGUUGU,<br>AAAUUUGGCUGAAAGAUGA |
| ATF4 | CCACGUUGGAUGACACUUGdTdT |

SI Methods

Visualising RNA/DNA hybrids by slot blot assay

The slot blot assay was adapted from a previously described method [1]. Cells were harvested in lysis buffer (100 mM Tris-HCl pH 8.5, 5 mM EDTA, 0.2% SDS, and 100 mM NaCl containing 0.5 mg/ml proteinase K) and incubated in a 55°C heated shaker overnight. Genomic DNA was extracted with isopropanol, washed in 70% ethanol, air-dried and re-suspended in TE buffer. 50 µg of genomic DNA was treated with 2 µL of RNase A (Fermentas) for 2 hours, and using a slot blot apparatus (Bio-Rad), equal amounts of samples were blotted onto a positively charged nylon transfer membrane (GE Healthcare). Where indicated, samples were treated with RNase H1 (NEB) overnight. The ssDNA blot was denatured (0.4 M NaOH and 0.6 M NaCl) and neutralised (1.5 M NaCl and 0.5 M Tris pH 7.4). Blots were UV crosslinked (0.12 $J/m^2$), blocked then incubated with S9.6 (1:1000, Kerafast) or ssDNA (1:5000, Millipore) antibodies and imaged using the Odyssey Infrared system.

RNA-sequencing Enrichment analysis

Several tools were used for the gathering of data on the differentially expressed genes. For G-quadruplex sequences, QuadBase was used [2]. We relied on BiomaRt for a series of measurements, namely gene length, number of exons and GC percent [3].

SI Figure Legends

Figure S1. **SETX is induced in response to hypoxia**

A. RKO cells were exposed to <0.1% $O_2$ for 0, 8 and 16 hours followed by western blotting with the indicated antibodies.

B. A549 cells were exposed to <0.1% $O_2$ for 0, 1.5, 6 and 12 hours with 5'EU (5-ethynyluridine) (0.5 mM) added for the final hour.

C. HCT116 cells were exposed to <0.1% $O_2$ for 6 hours followed by RT-qPCR for the relative mRNA levels of SETX determined by RT-qPCR.

D. HeLa cells were exposed to <0.1% $O_2$ for 6 hours followed by RT-qPCR for the relative mRNA levels of SETX determined by RT-qPCR.

E. U87-MG cells were exposed to <0.1% $O_2$ for 6 hours followed by RT-qPCR for the relative mRNA levels of SETX determined by RT-qPCR.

F. Expression analysis of SETX in The Cancer Genome Atlas (TCGA) datasets. RNA-sequencing data (RNA Seq V2 RSEM) for 382 colorectal adenocarcinoma tumours, 517 lung adenocarcinoma tumours and 501 lung squamous cell carcinoma tumours were downloaded from the TCGA project. A correlation of SETX was determined against hypoxia metagene signature [4]. Correlations and statistical significance were determined by calculating Spearman's rho rank correlation coefficient (r) and two-tailed *P* value using Hmisc package in RStudio.

(A-E) Data representative of two independent experiments (n=2). Mean ± standard error of the mean (SEM) are displayed.

Figure S2. **SETX depletion leads to increased R loops and replication stress in hypoxia**

A. RKO cells were exposed to 21% or <0.1% $O_2$ for 6 hours. Genomic DNA was extracted, treated with or without RNase H (RH) and subjected to slot blot assays with the S9.6 antibody. ssDNA was used as the loading control.

B. SETX transcript levels as determined by RT-qPCR in A549 cells with or without SETX siRNA after 16 h hypoxia (<0.1% $O_2$) treatment. Representative of 2 independent experiments.

C. Western blot analysis in A549 cells with or without SETX siRNA after 6 hours in normoxia or hypoxia (<0.1% $O_2$).

D. SETX transcript levels as determined by RT-qPCR in RKO cells with either siCtrl or SETX siRNA after 16 h hypoxia (<0.1% $O_2$) treatment.

E. Western blot analysis in RKO cells with control siRNA or SETX siRNA after 6 hours in normoxia or hypoxia (<0.1% $O_2$).

F. RKO cells were transfected with control siRNA (siCtrl) or SETX siRNA and exposed to 21% or <0.1% $O_2$ for 6 hours. Genomic DNA was extracted, treated with or without RNase H and subjected to slot blot assays with the S9.6 antibody. ssDNA was used as the loading control. There is a 3.5-fold increase in S9.6 signal in hypoxia, and a 2.3-fold increase in S9.6 signal upon SETX depletion.

G. Replication rate curves for RKO cells with control siRNA (siCtrl, grey) or SETX siRNA (green), as measured by IdU incorporation rates using immunofluorescence microscopy, in normoxia (21% $O_2$). Figure generated using the same data as Figure 2F.

H. Replication rate curves for RKO cells with control siRNA (siCtrl, grey) or SETX siRNA (green), as measured by IdU incorporation rates using immunofluorescence microscopy, after 6 hours of hypoxia (<0.1% $O_2$) treatment. Figure generated using the same data as Figure 2F.

(A-H) Data from three independent experiments (n=3), mean ± SEM are displayed unless otherwise indicated.

Figure S3. **SETX depletion affects transcription in hypoxia**

A. RKO control cells or SETX depleted cells were exposed to 21% and <0.1% $O_2$ for 6 hours in the presence of 5'EU. Cells were fixed, stained and quantified for 5'EU (0.5 mM) incorporation. n=1

B. Validation experiments of RNA-seq results. RKO cells were treated with control siRNA or SETX siRNA and exposed to <0.1% $O_2$ for 6 hours. RT-qPCR analysis was performed on a number of genes shown to be regulated by SETX in the RNA-seq data. For each gene, relative mRNA levels were determined using 18S as the reference gene. The fold change in expression upon SETX-depletion in hypoxia as compared to the control siRNA in hypoxia are shown. Genes in red were repressed in SETX depleted hypoxic cells, while genes in blue were induced, according to the RNA-seq results. Dotted line indicates no change in expression upon SETX depletion.

C. Gene ontology analysis found that a number of genes (69/341) upregulated in SETX depleted hypoxic cells were involved in rRNA processing, nucleolus and ribosome biogenesis.

(A-B) Data from three independent experiments (n=3), mean ± SEM are displayed unless otherwise indicated.

Figure S4. **Characteristics of genes affected by SETX depletion in hypoxia**

A. All genes from the RNA-sequencing were grouped into 3 sections: (1) Genes that do not change in hypoxia (column 1-3), (2) genes that are repressed in hypoxia (column 4-6), and (3) genes that are induced in hypoxia (column 7-9). Within each of the 3 sections, the length of genes was compared between genes do not change expression in siSETX hypoxic cells (green) to genes that decrease in siSETX hypoxic cells (yellow) and to genes that increase in siSETX hypoxic cells (orange).

B. Same analysis as A comparing number of exons.

C. Same analysis as A comparing number of G4 sequences.

D. Same analysis as A comparing GC content.

(A-D) Statistical significance was determined using the Wilcoxon test, and p-values were adjusted with the Holm method.

Figure S5. **SETX depletion and DNA damage in hypoxia.**

A. Percentage of cells displaying 53BP1 foci (>5 per nucleus) in RKO cells with either control siRNA or SETX siRNA. Cells were exposed to 21% $O_2$ (Norm) or <0.1% $O_2$ (Hyp) for 6 hours.

B. Percentage of cells displaying 53BP1 foci (>5 per nucleus) in RKO cells exposed to mock treatment, Adriamycin (ADR, 2 μM) or pre-treatment with DRB (100 μM) followed by Adriamycin (2 μM) for 6 hours. n=1

C. RKO cells were pre-treated with DMSO or DRB (100 μM) and exposed to 21% or <0.1% $O_2$ for 6 hours. Genomic DNA was extracted, treated with or without RNase H (RH) and subjected to slot blot assays with the S9.6 antibody. ssDNA was used as the loading control. The relative S9.6 signal normalised to the ssDNA control is shown on the left of the blot.

D. A549 control cells or cells treated with DRB (100 μM) for 6 hours, were fixed and stained with the S9.6 antibody and DAPI. Representative of 2 independent experiments.

E. Representative images from D.

F. RKO cells with mock or SETX siRNA were exposed to 21% $O_2$ (Norm) or 6 hours of <0.1% $O_2$ (Hyp) in the presence of BrdU (20 μM). Percentage of cells with 53BP1 foci (>5 per nucleus) that did not have BrdU incorporation (grey) or had BrdU incorporation (green) were determined. n=1

G. RKO cells with mock or SETX siRNA were exposed to 21% $O_2$ (Norm) or 6 hours of <0.1% $O_2$ (Hyp) in the presence of EdU (10 μM). Percentage of cells with 53BP1 foci (>5 per nucleus) that did not have EdU incorporation (grey) or had EdU incorporation (green) were determined. n=1

(A-G) Data from three independent experiments (n=3), mean ± SEM are displayed unless otherwise indicated.

Figure S6. **Hypoxia-induced SETX is independent of HIF1α or p53**

A. A549 cells were exposed to the oxygen concentrations shown for the times indicated followed by western blotting. Activation of PERK (electrophoretic mobility shift) and induction of GRP78 is indicative of activation of ATF3/4/5.

B. A549 cells were exposed to normoxia (21% $O_2$) and hypoxia (2% $O_2$ or < 0.1% $O_2$) for 16 hours, and CHOP mRNA levels were determined by RT-qPCR. Representative of 3 independent experiments.

C. A549 cells were treated with either control siRNA (grey) or HIF1α siRNA (green) and exposed to 21% $O_2$ (Norm) or <0.1% $O_2$ (Hyp) for 16 hours. SETX mRNA levels were determined by RT-qPCR.

D. A549 cells were treated with either control siRNA (grey) or HIF1α siRNA (green) and exposed to 21% $O_2$ (Norm) or <0.1% $O_2$ (Hyp) for 16 hours. VEGF mRNA levels were determined by RT-qPCR.

E. A549 cells were treated with either control siRNA (grey) or p53 siRNA (green) and exposed to 21 % $O_2$ (Norm) or <0.1% $O_2$ (Hyp) for 6 hours. PHLDA3 mRNA levels were determined by RT-qPCR.

(A-E) Data from three independent experiments (n=3), mean ± SEM are displayed unless otherwise indicated.

Figure S7. **Hypoxia-induced SETX is dependent on PERK/ATF4 signalling**

A. A549 cells were treated with either control siRNA (grey) or ATF4 siRNA (green) and exposed to 21% $O_2$ (Norm) or <0.1% $O_2$ (Hyp) for 16 hours. ATF4 mRNA levels were determined by RT-qPCR.

B. A549 cells were treated with either control siRNA (grey) or ATF4 siRNA (green) and exposed to 21% $O_2$ (Norm) or <0.1% $O_2$ (Hyp) for 16 hours. CHOP mRNA levels were determined by RT-qPCR.

C. A549 cells were exposed to 21% $O_2$ (Norm) or <0.1% $O_2$ (Hyp) for 4 hours, and ChIP-qPCR experiments were performed to determine fold enrichment of ATF4 at the CHOP promoter relative to IgG control.

D. CHOP mRNA levels in non-treated A549 cells (N) and cells exposed to tunicamycin (Tuni, 5 μg/mL) or thapsigargin (Thaps, 2 μM) for 16 hours.

E. Non-treated A549 cells (N), cells exposed to IR (5 Gy) followed by a 1 hour recovery, Hu (1 mM), DFO (100 μM) and hypoxia (2% $O_2$ or < 0.1% $O_2$) for 16 hours, were used in western blot assays to determine HIF1α, p53-S15, p53, γH2AX and H2AX protein levels.

F. A549 cells were exposed to Thaps (2 μM, 24 h) or Hu (2 mM, 8 h) and then stained for RPA. % of cells with more than 5 RPA foci are shown.

G. A549 cells were treated with either control siRNA (grey) or PERK siRNA (green) and exposed to 21% $O_2$ (Norm) or <0.1% $O_2$ (Hyp) for 16 hours. CHOP mRNA levels were determined by RT-qPCR. n=4

H. A549 cells were treated with either DMSO (grey) or PERK inhibitor (PERKi, 10 μM, green) and exposed to 21% $O_2$ (Norm) or <0.1% $O_2$ (Hyp) for 6 hours. CHOP mRNA levels were determined by RT-qPCR. n=4

I. A549 cells were pre-treated with or without PERKi (10 μM) and exposed to DMSO or Thaps (2 μM) for 6 hours. SETX mRNA levels were determined by RT-qPCR.

J. A549 cells were pre-treated with or without PERKi (10 μM) and exposed to DMSO or Thaps (2 μM) for 6 hours. CHOP mRNA levels were determined by RT-qPCR.

(A-J) Data from three independent experiments (n=3), mean ± SEM are displayed unless otherwise indicated.

Figure S8. **Loss of PERK leads to an accumulation of R-loops and replication stress**

A. A549 cells that were either mock transfected or transfected with PERK siRNA, were exposed to 21% $O_2$ or <0.1% $O_2$ for 6 hours. Cells were fixed and stained with the S9.6 antibody and DAPI. Where indicated, coverslips were treated with RNase H prior to staining. Representative images from Figure 5A. Dashed white line indicates nuclear outline based on DAPI stain that was used to measure the nuclear S9.6 intensity.

B. A549 cells that were treated with DMSO (grey) or PERKi (10 μM, green) were exposed to 21% $O_2$ or <0.1% $O_2$ for 12 hours. Cells were fixed and stained with the S9.6 antibody and DAPI. Where indicated, coverslips were treated with RNase H prior to staining.

C. A549 treated with DMSO or PERKi (10 μM) were exposed to 21% and <0.1% $O_2$ for 6 hours, 5'EU was added for the final hour of treatment. Cells were fixed, stained and quantified for 5'EU incorporation. n=1

D. A549 cells with either mock (grey) treatment or PERK siRNA (green) were exposed to hypoxia (<0.1% $O_2$) for 6 hours, cells were fixed and stained for RPA by immunofluorescence assays. Percentage of cells with greater than 5 RPA foci was quantified.

E. A549 cells treated with DMSO (grey) or PERKi (10 μM, green), were exposed to hypoxia (<0.1% $O_2$) for 6 hours, cells were fixed and stained for RPA by immunofluorescence assays. The number of RPA foci per cell was quantified.

(A-E) Data from two independent experiments (n=2), mean ± SEM are displayed unless otherwise indicated.

Figure S1

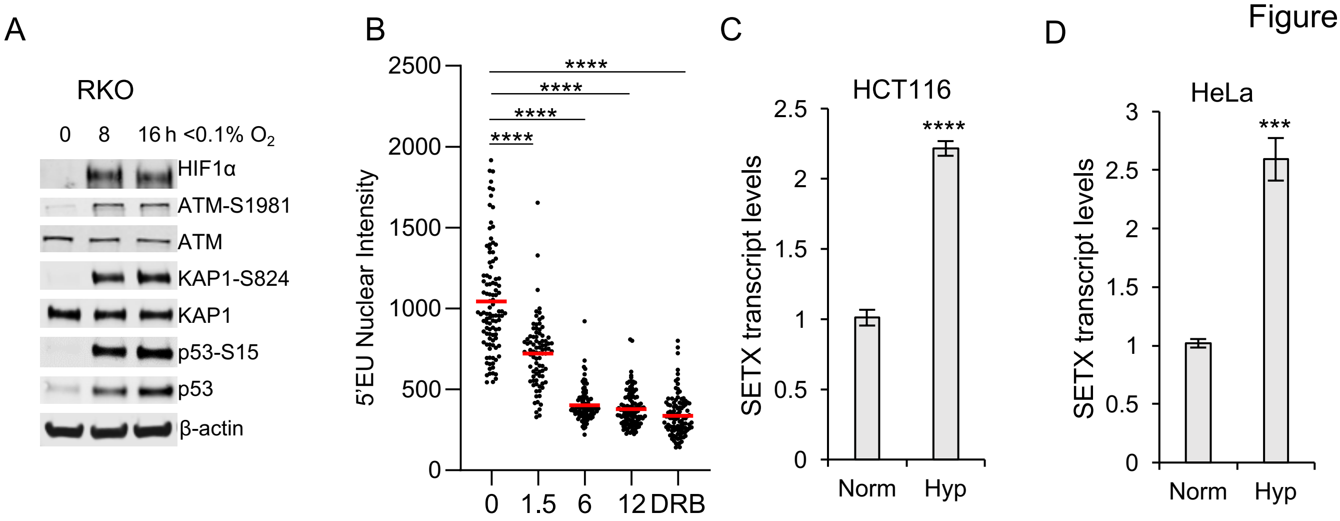


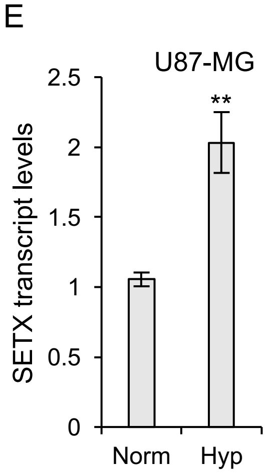


F

| Correlation of expression of SETX and hypoxia signature | | |
|---|---|---|
| **TCGA datasets:** | **Spearman:** | ***P*:** |
| Lung squamous cell carcinoma (n= 501) | -0.31 | <0.001 |
| Lung adenocarcinoma (n= 517) | -0.38 | <0.0001 |
| Colorectal (n=382) | -0.37 | <0.0001 |

Figure S2

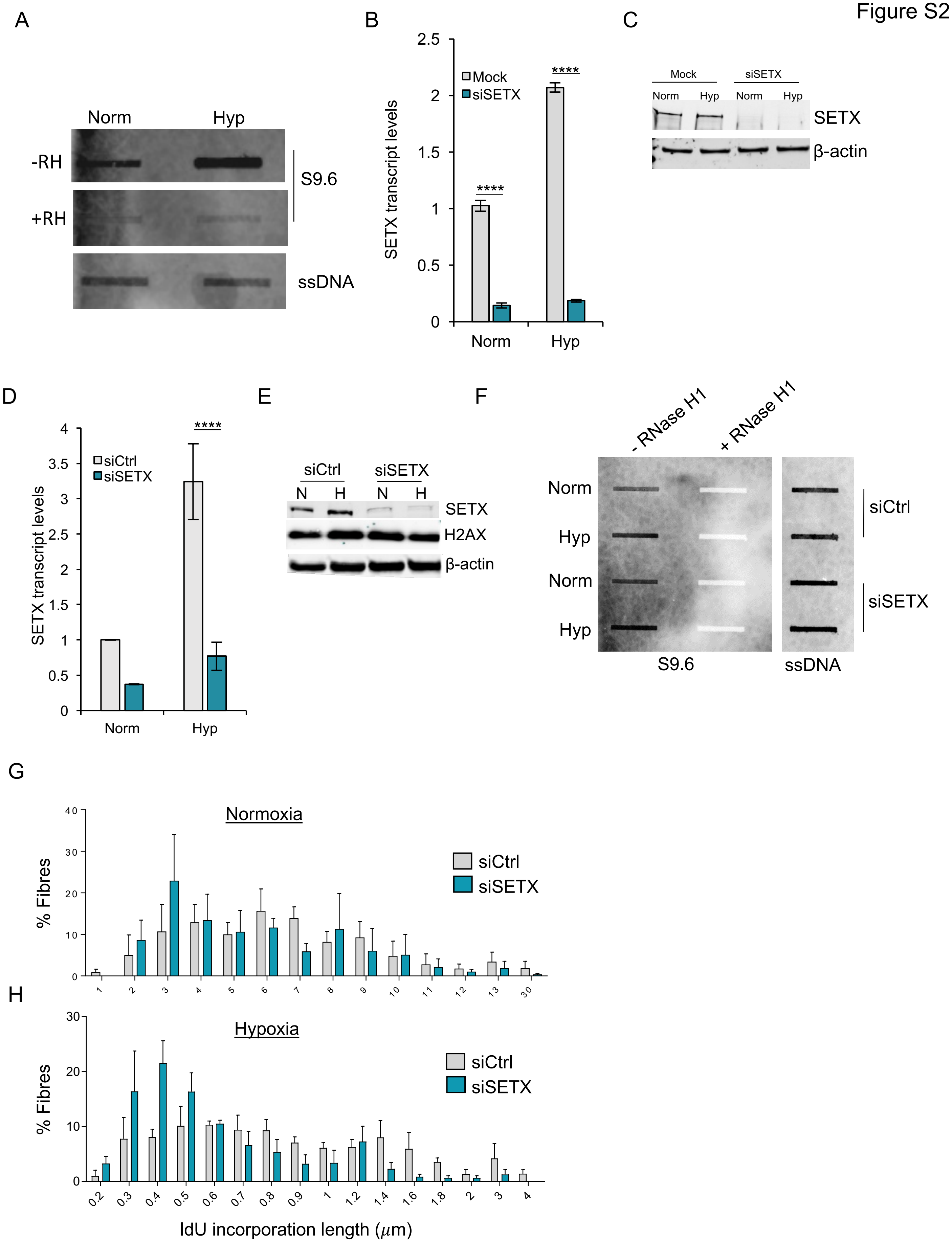
A
Norm
Hyp
-RH
+RH
S9.6
ssDNA
B
Mock
siSETX
SETX transcript levels
Norm
Hyp
****
****
C
Mock
siSETX
Norm
Hyp
Norm
Hyp
SETX
β-actin
D
siCtrl
siSETX
SETX transcript levels
Norm
Hyp
****
E
siCtrl
siSETX
N
H
N
H
SETX
H2AX
β-actin
F
- RNase H1
+ RNase H1
Norm
Hyp
Norm
Hyp
siCtrl
siSETX
S9.6
ssDNA
G
Normoxia
siCtrl
siSETX
% Fibres
H
Hypoxia
siCtrl
siSETX
% Fibres
IdU incorporation length (μm)

### Figure S3

A

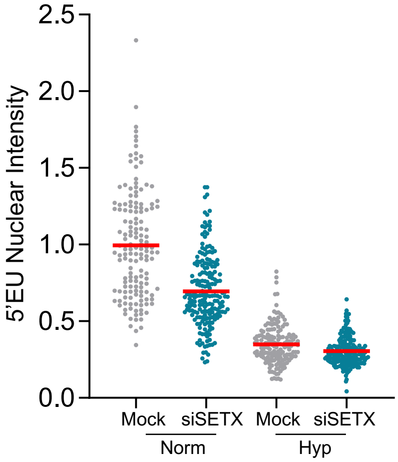


B

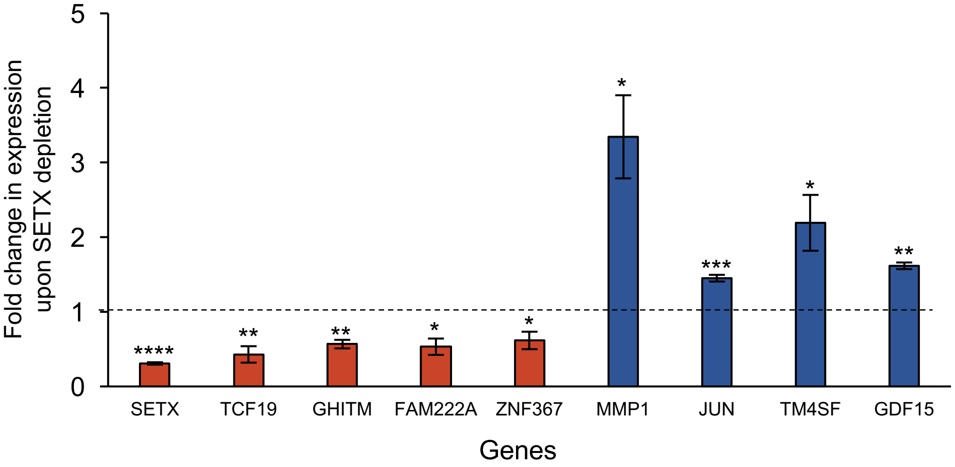


C

| term ID | term name |
|---|---|
| GO:0034660 | ncRNA metabolic process |
| GO:0016072 | rRNA metabolic process |
| GO:0022613 | ribonucleoprotein complex biogenesis |
| GO:0042254 | ribosome biogenesis |
| GO:0034470 | ncRNA processing |
| GO:0006364 | rRNA processing |
| GO:0030490 | maturation of SSU-rRNA |
| GO:0030684 | preribosome |
| GO:0032040 | small-subunit processome |
| GO:0005730 | nucleolus |
| GO:0044452 | nucleolar part |
| GO:0030515 | snoRNA binding |
| CORUM:5101 | CyclinD3-CDK4-CDK6-p21 complex |

Figure S4

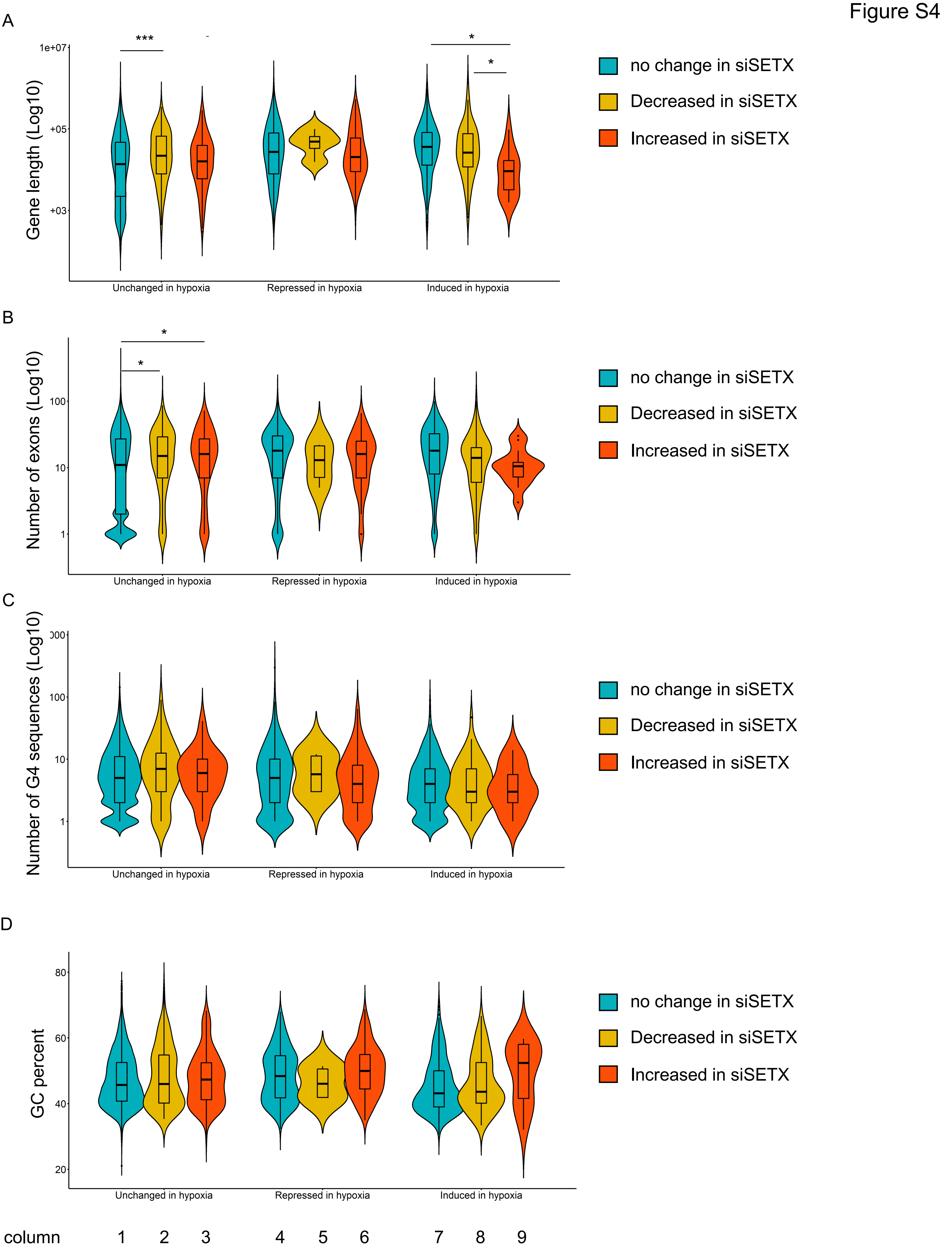
A
***
*
*
1e+07
+05
+03
Gene length (Log10)
Unchanged in hypoxia
Repressed in hypoxia
Induced in hypoxia
no change in siSETX
Decreased in siSETX
Increased in siSETX
B
*
*
100
10
1
Number of exons (Log10)
Unchanged in hypoxia
Repressed in hypoxia
Induced in hypoxia
no change in siSETX
Decreased in siSETX
Increased in siSETX
C
100
10
1
Number of G4 sequences (Log10)
Unchanged in hypoxia
Repressed in hypoxia
Induced in hypoxia
no change in siSETX
Decreased in siSETX
Increased in siSETX
D
80
60
40
20
GC percent
Unchanged in hypoxia
Repressed in hypoxia
Induced in hypoxia
no change in siSETX
Decreased in siSETX
Increased in siSETX
column
1
2
3
4
5
6
7
8
9

Figure S5

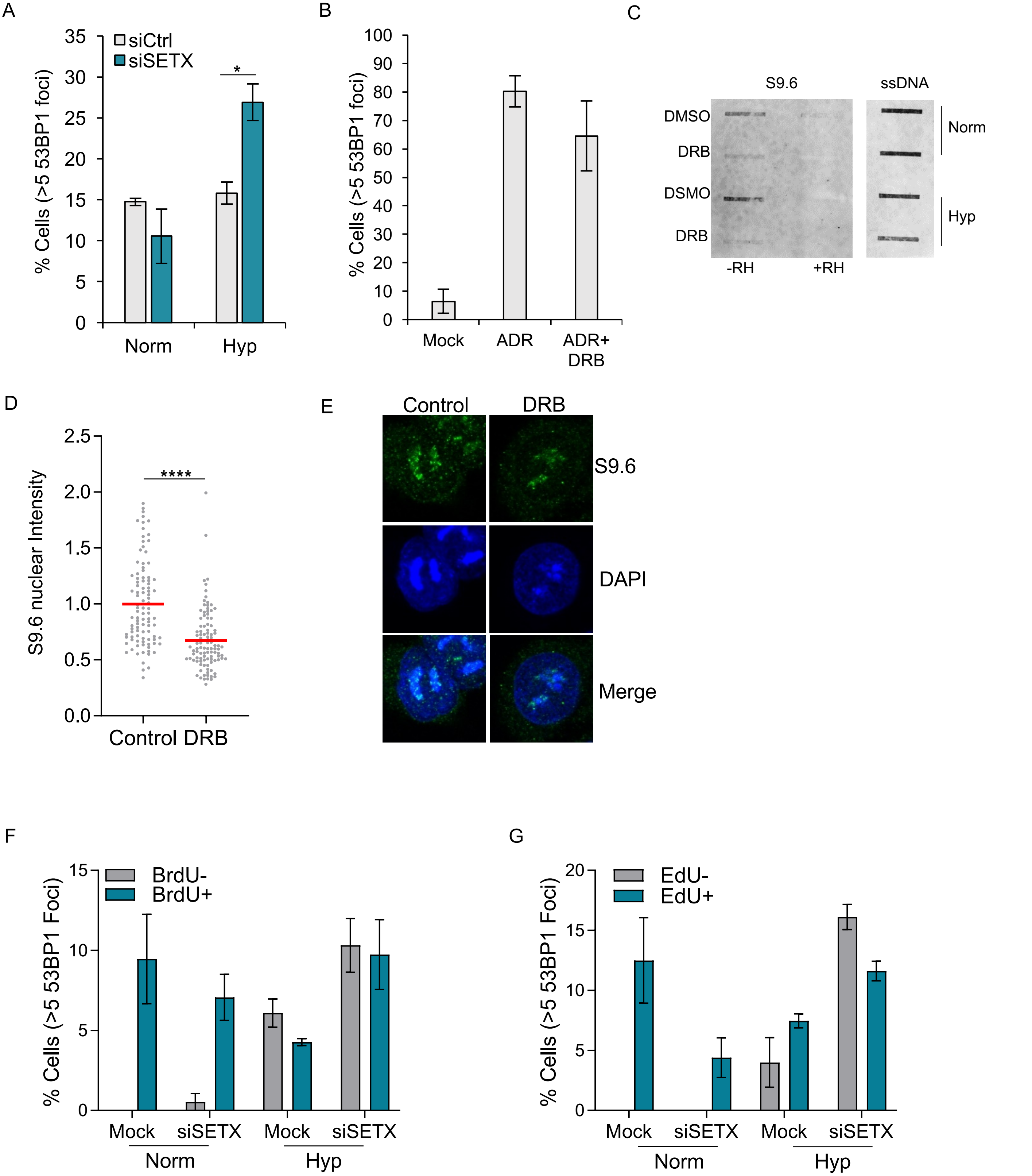
A
% Cells (>5 53BP1 foci)
siCtrl
siSETX
*
Norm
Hyp
B
% Cells (>5 53BP1 foci)
Mock
ADR
ADR+ DRB
C
S9.6
ssDNA
DMSO
DRB
DSMO
DRB
Norm
Hyp
-RH
+RH
D
S9.6 nuclear Intensity
****
Control DRB
E
Control
DRB
S9.6
DAPI
Merge
F
% Cells (>5 53BP1 Foci)
BrdU-
BrdU+
Mock
siSETX
Mock
siSETX
Norm
Hyp
G
% Cells (>5 53BP1 Foci)
EdU-
EdU+
Mock
siSETX
Mock
siSETX
Norm
Hyp

Figure S6

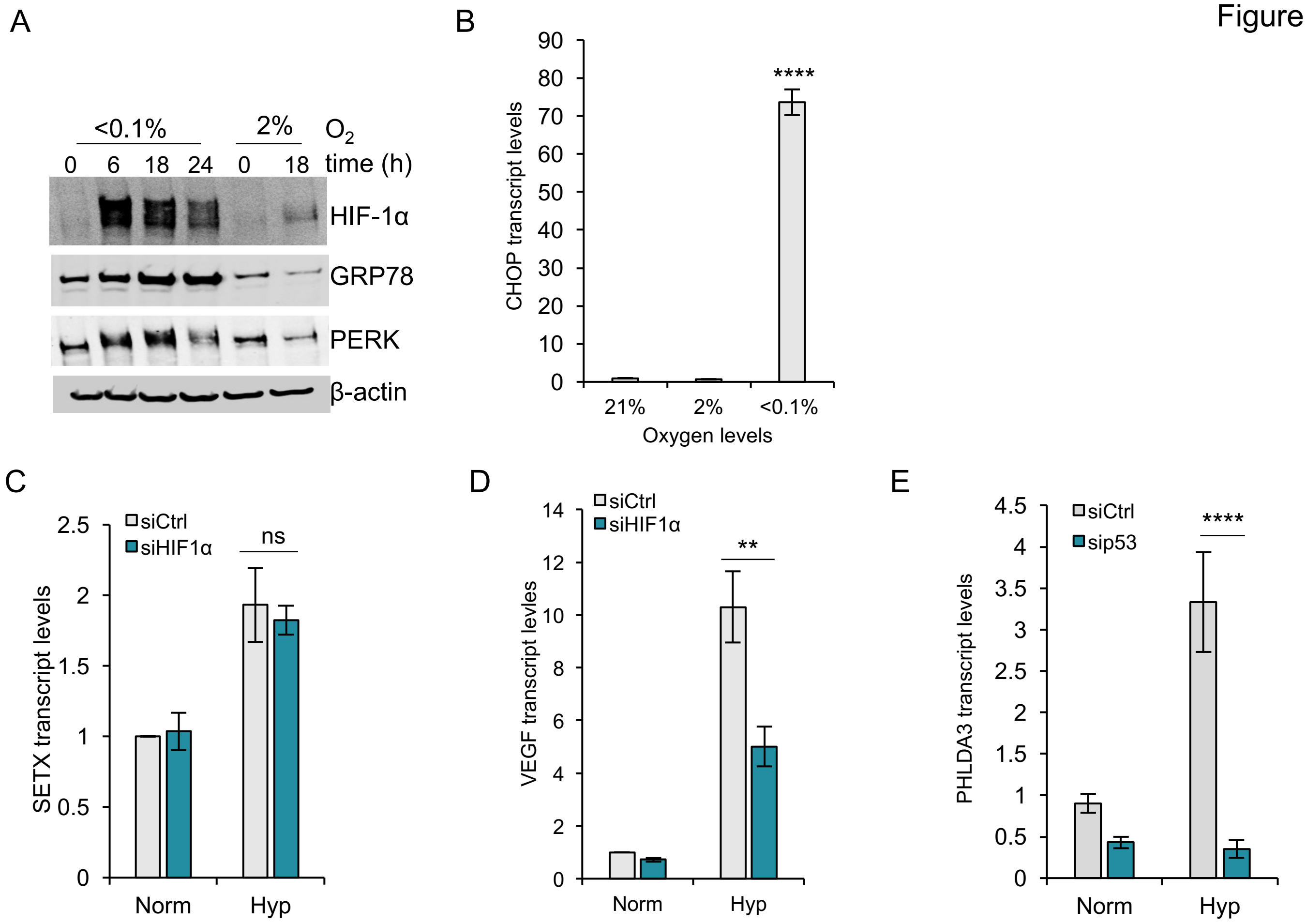
A
<0.1%
2%
$O_2$
0 6 18 24 0 18 time (h)
HIF-1α
GRP78
PERK
β-actin
B
CHOP transcript levels
90
80
70
60
50
40
30
20
10
0
****
21%
2%
<0.1%
Oxygen levels
C
SETX transcript levels
2.5
2
1.5
1
0.5
0
siCtrl
siHIF1α
ns
Norm
Hyp
D
VEGF transcript levles
14
12
10
8
6
4
2
0
siCtrl
siHIF1α
**
Norm
Hyp
E
PHLDA3 transcript levels
4.5
4
3.5
3
2.5
2
1.5
1
0.5
0
siCtrl
sip53
****
Norm
Hyp

Figure S7

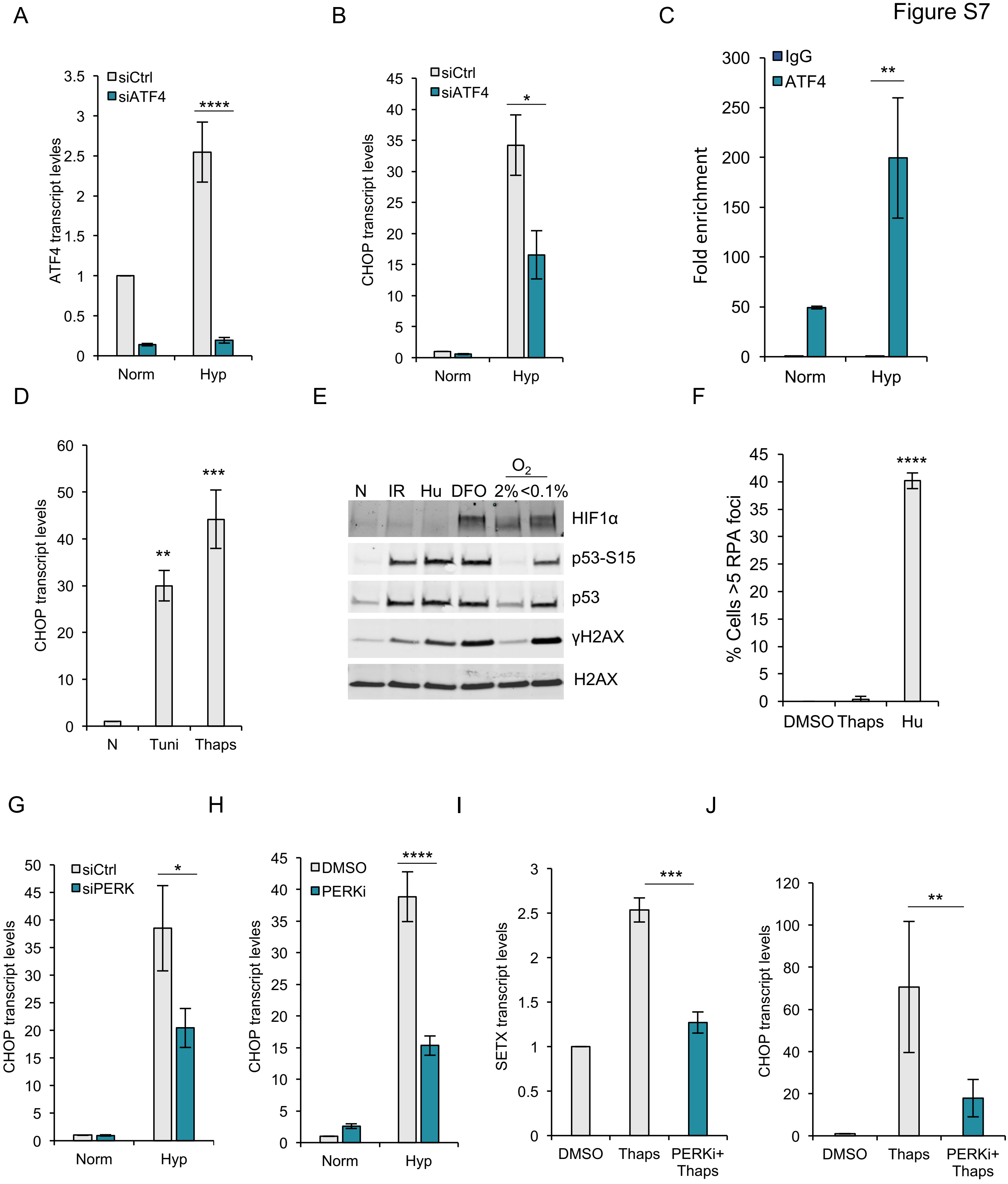
A
B
C
D
E
F
G
H
I
J
ATF4 transcript levles
siCtrl
siATF4
****
Norm
Hyp
CHOP transcript levels
*
Fold enrichment
IgG
ATF4
**
N
Tuni
Thaps
***
$O_2$
N IR Hu DFO 2% <0.1%
HIF1α
p53-S15
p53
γH2AX
H2AX
% Cells >5 RPA foci
DMSO
Thaps
Hu
siPERK
CHOP transcript levles
PERKi
SETX transcript levels
PERKi+
Thaps

### Figure S8

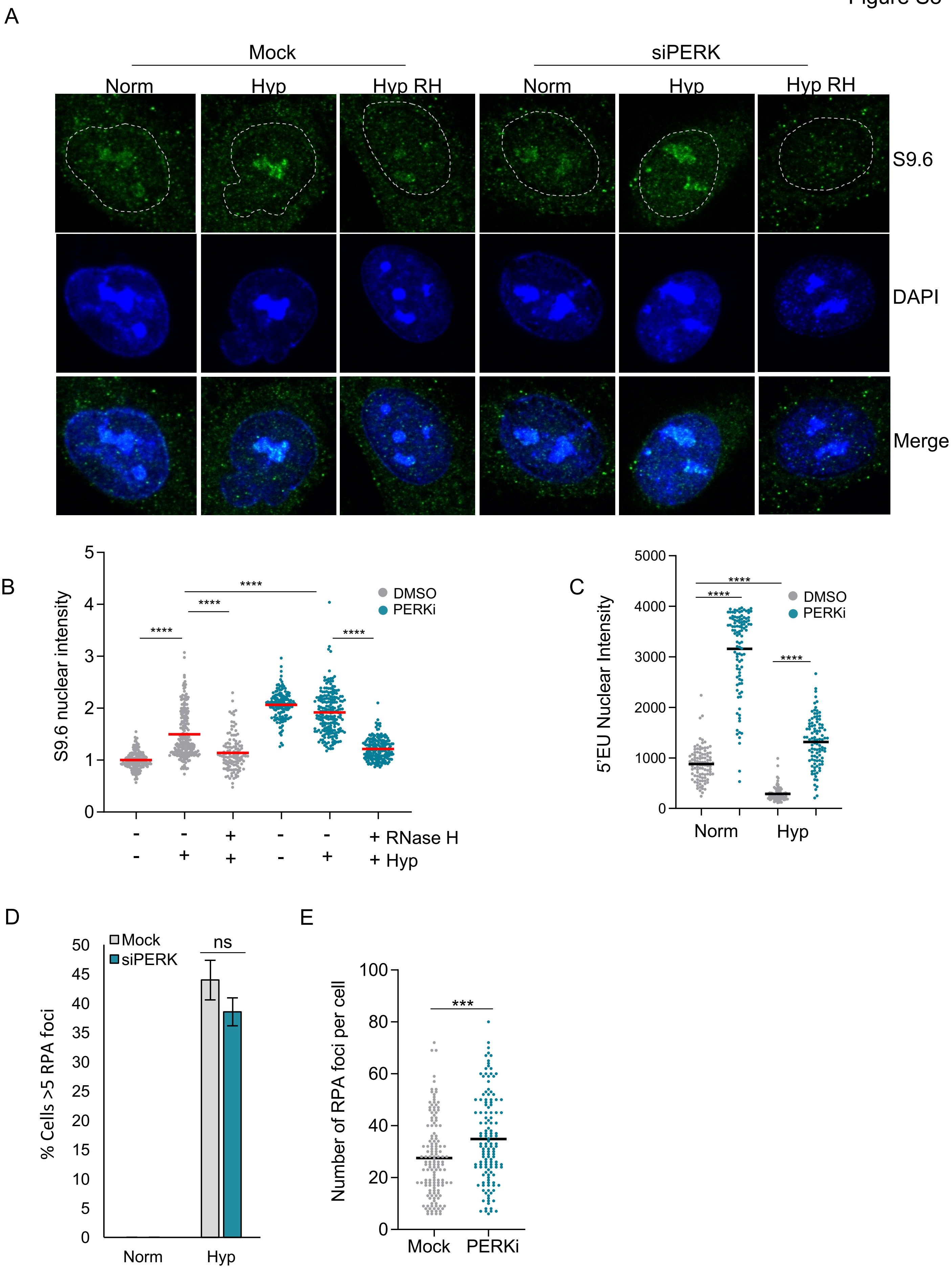
A
Mock
siPERK
Norm
Hyp
Hyp RH
Norm
Hyp
Hyp RH
S9.6
DAPI
Merge
B
S9.6 nuclear intensity
DMSO
PERKi
****
****
****
****
- - + - - + RNase H
- + + - + + Hyp
C
5'EU Nuclear Intensity
DMSO
PERKi
****
****
****
Norm
Hyp
D
% Cells >5 RPA foci
Mock
siPERK
ns
Norm
Hyp
E
Number of RPA foci per cell
***
Mock
PERKi
